# Isoform-specific C-terminal phosphorylation drives autoinhibition of Casein Kinase 1

**DOI:** 10.1101/2023.04.24.538174

**Authors:** Rachel L. Harold, Nikhil K. Tulsian, Rajesh Narasimamurthy, Noelle Yaitanes, Maria G. Ayala Hernandez, Hsiau-Wei Lee, Priya Crosby, Sarvind M. Tripathi, David M. Virshup, Carrie L. Partch

**Author notes:** Equal contributions (Rachel L. Harold, Nikhil K. Tulsian).

## Abstract

Casein kinase 1 δ (CK1δ) controls essential biological processes including circadian rhythms and Wnt signaling, but how its activity is regulated is not well understood. CK1δ is inhibited by autophosphorylation of its intrinsically disordered C-terminal tail. Two CK1 splice variants, δ1 and δ2, are known to have very different effects on circadian rhythms. These variants differ only in the last 16 residues of the tail, referred to as the extreme C-termini (XCT), but with marked changes in potential phosphorylation sites. Here we test if the XCT of these variants have different effects in autoinhibition of the kinase. Using NMR and HDX-MS, we show that the δ1 XCT is preferentially phosphorylated by the kinase and the δ1 tail makes more extensive interactions across the kinase domain. Mutation of δ1-specific XCT phosphorylation sites increases kinase activity both *in vitro* and in cells and leads to changes in circadian period, similar to what is reported *in vivo*. Mechanistically, loss of the phosphorylation sites in XCT disrupts tail interaction with the kinase domain. δ1 autoinhibition relies on conserved anion binding sites around the CK1 active site, demonstrating a common mode of product inhibition of CK1δ. These findings demonstrate how a phosphorylation cycle controls the activity of this essential kinase.

**Significance:** Subtle control of kinase activity is critical to physiologic modulation of multiple physiological processes including circadian rhythms. CK1δ and the closely related CK1ε regulate circadian rhythms by phosphorylation of PER2, but how kinase activity itself is controlled is not clear. Building on the prior observation that two splice isoforms of CK1δ regulate the clock differently, we show that the difference maps to three phosphorylation sites in the variably spliced region (XCT) that cause feedback inhibition of the kinase domain. More broadly, the data suggest a general model where CK1 activity on diverse substrates can be controlled by signaling pathways that alter tail phosphorylation. These inhibitory phosphorylation sites could also be targets for new therapeutic interventions.

## Introduction

Kinases play a crucial role in biology through their ability to post-translationally modify proteins by phosphorylation, altering the structure and/or activity of their targets. The activity of most kinases is therefore kept under tight regulation, requiring phosphorylation of their activation loops and/or binding to cofactors or scaffolds to remodel the active site and recruit substrates. In some cases, removal of inhibitory phosphorylation can activate kinases (1, 2). The kinase domains of Casein Kinase 1δ (CK1δ) and its paralog, CK1ε, are constitutively active, but autophosphorylation of their intrinsically disordered C-terminal extensions, or ‘tails’, inhibits kinase activity (3, 4). CK1δ/ε tails are also targeted by a number of other kinases (5, 6) and phosphatases (4, 7–9) *in vivo,* indicating a dynamic mode of kinase regulation that is not well understood. Given the involvement of CK1δ/ε in a wide range of pathways from Wnt signaling, cell division, and apoptosis to circadian rhythms (5), understanding how the kinase activity of CK1δ/ε is regulated by phosphorylation of their intrinsically disordered tails could shed light on underappreciated regulatory mechanisms.

In mammalian circadian rhythms, CK1δ/ε regulate the intrinsic timing of the molecular clock through phosphorylation of the Period (PER) proteins (10–12). A phosphoswitch involving competing CK1δ/ε-dependent sites on PER2 regulates its half-life (13), wherein phosphorylation of β-TrCP degrons and subsequent protein turnover are counteracted by phosphorylation of several serines located within the Casein Kinase 1 binding domain (CK1BD) that anchors the kinase to PER throughout its daily life cycle (14–17). These stabilizing phosphorylation sites occur within the Familial Advanced Sleep Phase (FASP) region, named for a point mutation in human PER2 (S662G) that eliminates the priming phosphorylation and shortens the clock by ∼4 hours (18, 19). We recently showed that phosphorylation of the PER2 FASP region leads to feedback inhibition of the anchored kinase, utilizing two highly conserved anion binding sites on CK1δ near the active site that bind phosphoserines in the FASP to block access to substrates (20). This is similar to feedback inhibition of the kinase observed in DNA damage-induced apoptosis (21). Conservation of these feedback inhibition binding modes establishes the importance of CK1δ/ε anion binding sites for regulation of the kinase by phosphorylated substrates.

Seemingly modest alterations in CK1δ/ε activity are well established to shift the balance of the phosphoswitch to have a profound effect on circadian timing in mammals from rodents to humans. These alterations include inherited polymorphisms (22–25) and alternative splicing of CK1δ (26). The two CK1δ splice isoforms, δ1 and δ2, are identical throughout the kinase domain and most of the intrinsically disordered tail, differing only in the last 16 amino acids of the C-terminus (Fig. 1A), a region we call here the extreme C-terminus, or XCT. Despite this relatively minor change, the δ1 and δ2 isoforms differentially phosphorylate the PER2 FASP region *in vivo* to control circadian period (26). Truncating the XCT at residue 400 is sufficient to derepress kinase activity (27), suggesting a crucial role for the XCT in regulation of CK1 activity in cells.

**Figure 1.**
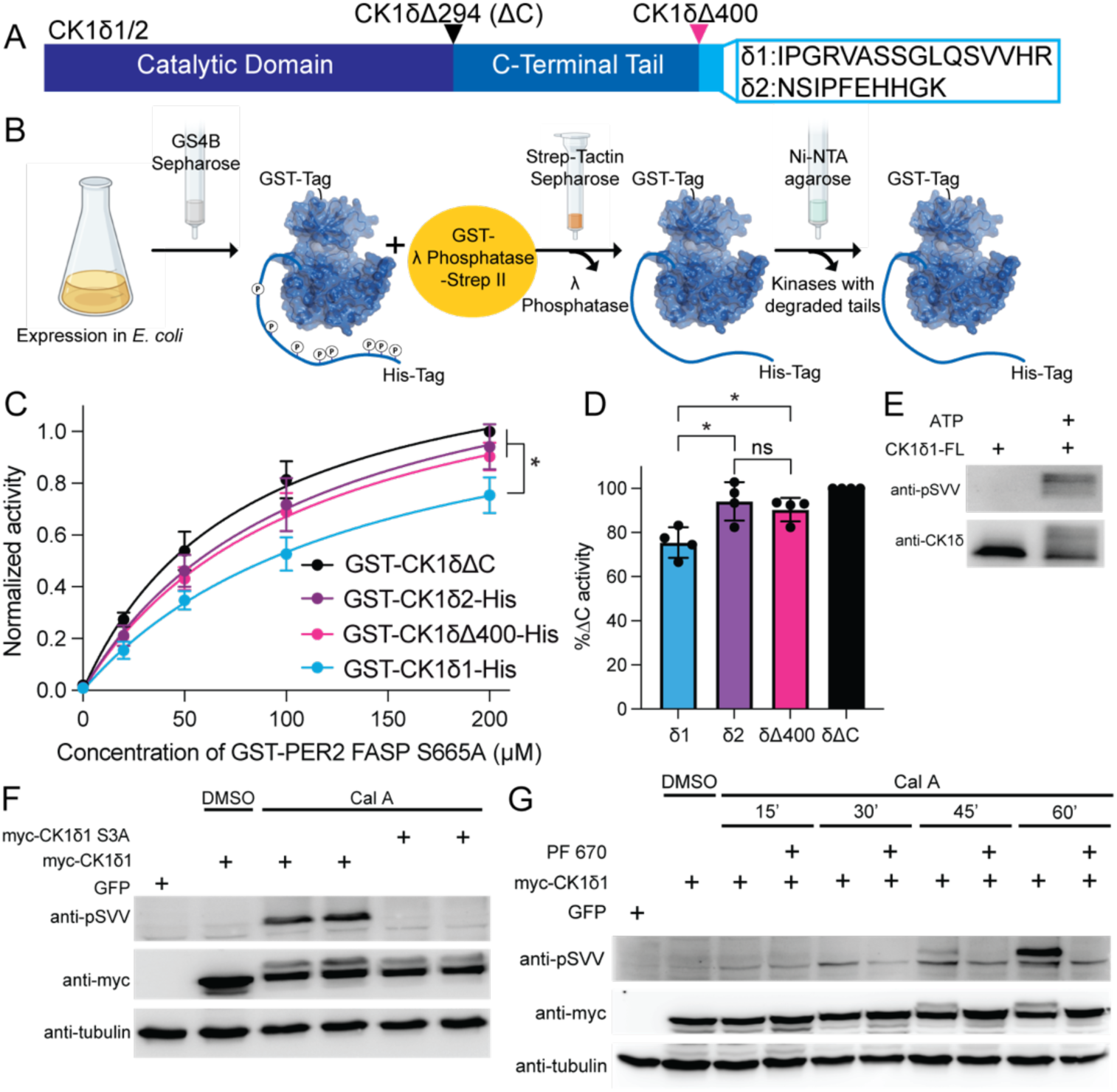
CK1δ isoforms change kinase activity *in-vitro* and regulation of XCT phosphorylation in cells. **A**, Schematic of CK1δ isoforms and truncations. **B**, Kinase purification scheme with double tag system. **C**, Quantification of ^32^P kinase assay measuring priming phosphorylation by CK1δ1 (blue), CK1δ2 (purple), CK1δΔ400 (pink), or CK1δΔC (black), normalized to max activity of CK1δΔC. Significance assessed by Extra sum-of-squares F Test (n=4, mean ± s.d.) **D**, Quantification of endpoint (200μM) in panel C. Ordinary one-way ANOVA, *, p<0.05 (n=4, mean ± s.d.). **E,** Representative western blot of purified CK1δ1 autophosphorylation detected with pSVV ab (n=3). **F,** Representative western blot of myc-CK1δ1 or myc-CK1δ1 S3A (S406/S407/S411A) phosphorylation in cells treated with calyculin A (n=3). **G,** Representative western blot of CK1δ1 phosphorylation in cells treated with Cal A and PF 670 (n=3).

In this study, to understand how the splice variants differentially regulate clock speed, we examined the biochemical role of the XCT in the regulation of CK1δ isoform activity. We found that isoform-specific differences in CK1 activity are intrinsic to the kinase *in vitro*; CK1δ1 is autoinhibited to a greater extent than CK1δ2, and truncation of the isoform-specific extreme C-termini essentially eliminates autoinhibition. NMR assays of the δ1 tail with the kinase domain in *trans* or *cis*, via segmental isotopic labeling, revealed that interaction between the kinase domain and δ1 tail is dependent on phosphorylation and predominantly localized to the δ1-specific sequence. By contrast, the δ2 tail interacts with the kinase domain even in the absence of phosphorylation, consistent with inhibition data using isolated δ1 or δ2 peptides. Biochemically, mutation of three phosphosites in the unique region of CK1δ1 increased its activity to that of CK1δ2. Using hydrogen/deuterium exchange mass spectrometry (HDX-MS), we mapped how CK1 isoforms or mutants differentially protect the kinase domain and intrinsically disordered tails from deuterium exchange. Protection of the kinase active site in the presence of an inhibitory phosphoFASP peptide of PER2 (20) mirrors protection by the two CK1δ isoforms, which is largely eliminated by deleting the XCT region with the Δ400 truncation or mutation of the three δ1-specific phosphosites. These biochemical findings translate in cells, where mutation of the δ1-specific phosphosites increases kinase activity and effects the period of the cell’s circadian rhythm. Our study defines isoform-specific mechanisms of CK1δ regulation and demonstrates a conserved mechanism of kinase inhibition by phosphorylated substrates.

## Results

### CK1δ isoforms δ1 and δ2 have different kinase activity *in vitro*

The alternatively spliced CK1δ isoforms, δ1 and δ2, differ only in the last 16 amino acids of the C-terminal tail (Fig. 1A). We previously showed that CK1δ1, CK1δ2, and a related isoform, CK1ε, can all catalyze the rate-limiting priming phosphorylation of PER2 FASP, with CK1δ2 and CK1ε demonstrating increased activity compared to CK1δ1 (27). This is biologically relevant, as CK1δ1 expression causes a shorter circadian period than CK1δ2 (26). The kinase domains of CK1δ and CK1ε are highly conserved, and though their intrinsically disordered C-terminal tails differ, the XCT of CK1δ2 and CK1ε are quite similar but dissimilar to the XCT of CK1δ1 (Fig. S1A). This suggested that the differences in biological activity we observed in cells might be due to their XCT; consistent with this, truncation of the XCT of CK1δ1 (Δ400) eliminated its isoform-specific decrease in activity (27).

To determine if isoform-specific differences in activity are intrinsic to the kinase, we established a pipeline to purify dephosphorylated full-length recombinant proteins (Fig. 1B). To demonstrate the effect of tail phosphorylation on CK1δ1, we first pre-treated full-length dephosphorylated CK1δ1 with ATP to induce autophosphorylation and then measured its activity on the PER2 FASP priming site. As expected, pretreatment of the kinase with ATP before the substrate was added significantly reduced FASP priming phosphorylation compared to untreated CK1δ1 (Fig. S1B). We then used *in vitro* kinase assays to measure the activity of the two CK1δ isoforms compared to the Δ400 truncation and the fully active, tail-less CK1δΔC. Activity was measured on a S665A mutant of the human PER2 FASP residues 645–687 fused to GST (Fig. 1C and S1C). The S665A mutation only allows for phosphorylation of the rate-limiting priming serine at S662 (20). Although differences in activity were modest, the activity of CK1δ1 on the PER2 FASP was significantly lower than CK1δ2 (Fig. 1C, D) and consistent with observed biological differences between CK1δ1 and CK1δ2 (26, 27). Truncation of the XCT or the entire tail also gave rise to activity that was similar to CK1δ2, recapitulating differences in the activity we observed in cells (27) and suggesting that CK1δ1 has increased autoinhibition. Tracking autophosphorylation with kinase assays (Fig. S1C, D) and Coomassie gel shift (Fig. S1G, H) showed that relative to the full-length proteins, the isolated kinase domain showed minimal autophosphorylation, and the δ1 kinase was significantly less phosphorylated than δ2 or Δ400, despite having more serines (Fig. S1D, H). These differences point to a potential regulatory role for the CK1δ1 XCT.

### Regulation of XCT autophosphorylation by phosphatases in cells

To test if any of the XCT phosphorylation sites are indeed autophosphorylated in cells, we used an antibody that specifically recognizes a phosphorylated serine in the sequence SVV (pSVV antibody) that corresponds to S411 of CK1δ1 (Fig. S1I, J). This antibody detected robust autophosphorylation of purified full-length CK1δ1 (Fig. 1E). We then assessed phosphorylation of the XCT region in cells expressing myc-tagged CK1δ1. Phosphorylation of the CK1δ tail is regulated *in vivo* by an active cycle of autophosphorylation and dephosphorylation (7). Consistent with this, accumulation of S411 phosphorylation was only detected when cells were treated with calyculin A, a broad-spectrum inhibitor of cellular Ser/Thr phosphatases, revealing reversible phosphorylation of CK1δ1 S411 in cells (Fig 1F). This is also consistent with reports of CK1δ1 phosphorylation at residues S406, S407, and S411 from large-scale phosphoproteomics screens (28, 29). Phosphorylation of CK1δ1 S411 in cells was blocked by addition of the CK1δ/ε-specific inhibitor PF6700462 (PF670) (Fig 1G), indicating that it is primarily due to autophosphorylation. These data establish that S411 in the XCT of CK1δ1 is autophosphorylated in cells in a reversible and dynamic manner.

Our *in vitro* results clearly indicate that the length and phosphorylation status of CK1δ isoforms influences their kinase activity. Moreover, the finding of autophosphorylation and phosphatase-mediated dephosphorylation of the XCT region in cells demonstrates the potential for dynamic regulation of CK1δ1 activity *in vivo*. Multiple studies have reported regulation of CK1δ kinase activity from structural and biochemical perspectives, including examining the role of autophosphorylation of the tail region (12, 20–22, 26, 27, 30). These studies have not addressed how the recently described splice isoforms of CK1δ can regulate its kinase activity. Accordingly, we next employed NMR and HDX-MS to develop mechanistic insights into this process.

### Differences in regulation of the δ1 and δ2 tails by the kinase domain

Possible phosphorylation sites in the tail have been mapped on full-length CK1δ1 *in vitro* (3) and in cells (31), where a non-phosphorylatable (NP) mutant substituting all 28 serines and threonines in the tail to alanine eliminated its electrophoretic mobility shift and regulation by cellular protein phosphatases (6). However, these studies did not unambiguously identify functionally important phosphorylation sites nor regions required for autoinhibition. The δ1 tail is 125 amino acids long and predicted to be intrinsically disordered, and no part of it was captured in a crystal structure of the full-length kinase (32). Therefore, we turned to nuclear magnetic resonance (NMR) spectroscopy to monitor backbone chemical shifts on the ^15^N δ1 tail alone or in the presence of equimolar amounts of the unlabeled kinase domain to better understand potential regulatory mechanisms. The ^15^N-^1^H HSQC spectrum of the isolated ^15^N δ1 tail exhibited limited chemical shift dispersion consistent with an intrinsically disordered protein, and addition of kinase domain in the absence of ATP resulted in very minor chemical shift perturbations (Fig. 2A, left), indicating that the tail and kinase domain did not interact in *trans* at these concentrations. However, incubation of the same sample with ATP for 2 hours at 25 °C resulted in chemical shift perturbations in the ^15^N δ1 tail backbone, as well as several new peaks downfield that correspond to phosphorylated residues (Fig. 2A, right), demonstrating that the kinase can phosphorylate its tail in *trans*.

**Figure 2.**
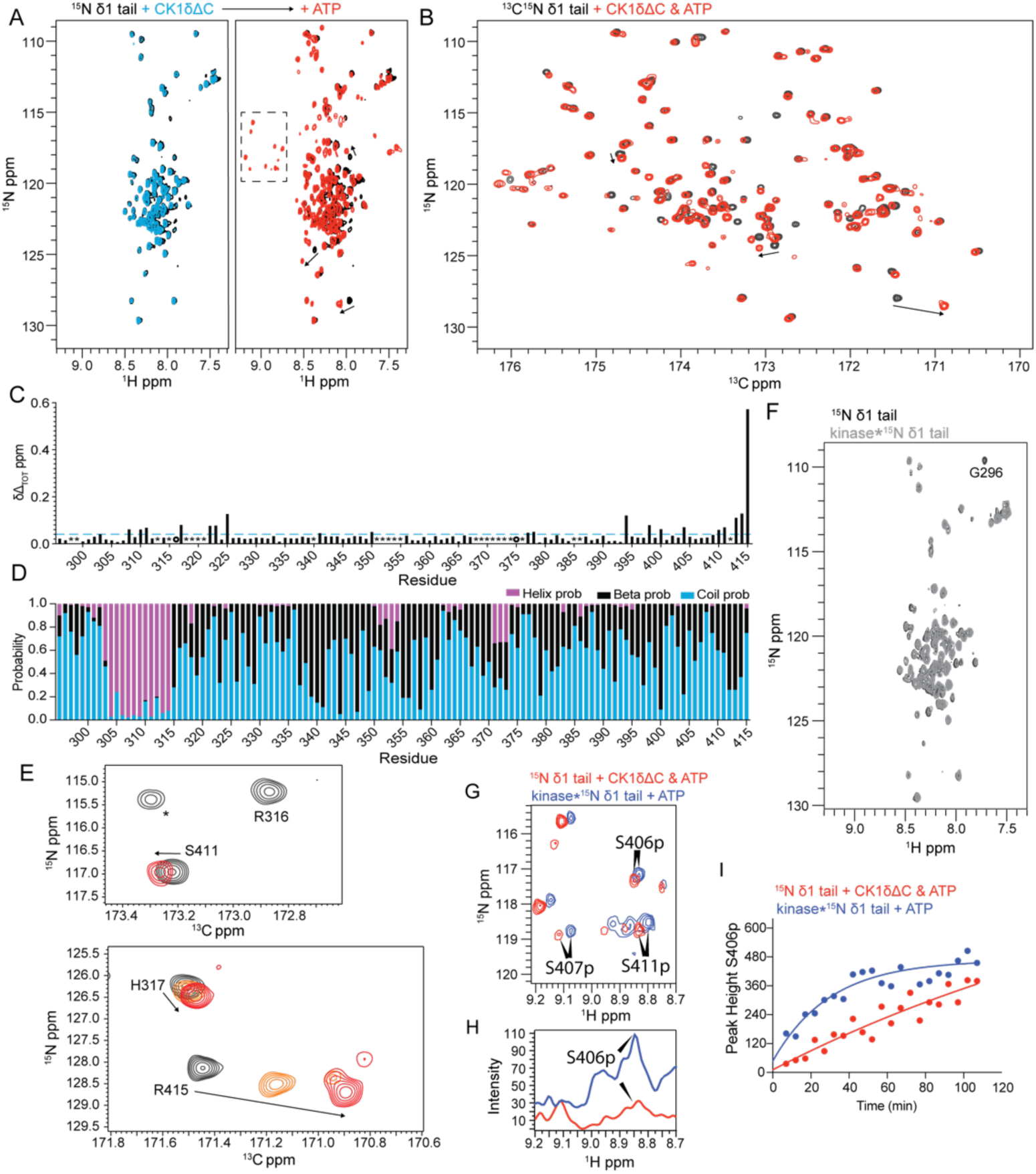
Phosphorylation-dependent interaction of the δ1 tail with the kinase domain. **A**, ^15^N-^1^H HSQC spectra of 100 μM ^15^N δ1 tail alone (black) or in the presence of 100 μM CK1δΔC (blue), or 100 μM CK1δΔC and 2.5 mM ATP for 2 hours at 25 °C (red); dashed box, peaks corresponding to phosphorylated residues. Arrows, chemical shift perturbation with phosphorylation. **B**, ^15^N-^13^C CON spectra of 100 μM ^13^C,^15^N δ1 tail alone (black) or in the presence of 100 μM CK1δΔC and 2.5 mM ATP for 2 hours at 25°C (red). Arrows, chemical shift perturbation of the same peaks from panel B. **C**, Quantification of chemical shift perturbations (Δδ_TOT_, ppm) from the ^15^N-^13^C CON spectrum of the δ1 tail alone or after incubation with CK1δΔC and ATP. Dashed line, significance cutoff at 0.05 ppm; open circle, broadened peaks; *, unassigned peaks. **D**, Secondary Structure Propensity (SSP) analysis of the δ1 tail from chemical shift data. Random coil (blue), ɑ-helix (pink), β-strand (black). **E**, Zoomed in view of CON spectra showing phosphorylation-dependent changes in the extreme C-terminus and predicted ɑ-helical region. δ1 tail alone (black), with CK1δΔC and ATP for 2 hours at 25 °C (red), or an unquenched partial phosphorylation reaction (orange). **F,** ^15^N-^1^H HSQC spectra of 34 μM δ1 tail alone (black) or in segmentally labeled kinase (gray). G296, only residue broadened in full-length kinase. **G**, ^15^N-^1^H HSQC spectra of ^15^N δ1 tail with CK1δΔC and ATP in *trans* (red) or in *cis* (blue) with phosphopeak assignments. **H**, ^15^N-edited 1D ^1^H spectra of phosphopeaks in panel G. **I**, Quantification of 1D time course measuring phosphorylation of ^15^N δ1 tail S406 in *trans* (red) or in *cis* (blue) with kinase.

Due to issues with peak overlap in the ^15^N-^1^H HSQC spectra, we turned to ^13^C-direct detection experiments, using ^15^N-^13^C CON spectra that provide better resolution of the protein backbone in disordered proteins (33). Addition of unlabeled kinase to the ^13^C,^15^N δ1 tail in the absence of ATP showed minimal chemical shift perturbation in the tail, similar to our ^15^N-^1^H HSQC data (Fig. S2A). After ATP addition, we observed chemical shift perturbations at the same resonances as in the HSQC (Fig. 2B), as well as some peak broadening upstream of XCT. Shifts were localized primarily to the XCT of the δ1 tail, and to a lesser extent in a charged region from residues 305-320 that is predicted to have alpha-helical structure (Fig. 2C-E). Peak broadening and chemical shift perturbations indicate that the XCT as well as some upstream residues are interacting with the kinase domain dependent on autophosphorylation. By contrast, ^15^N-^1^H HSQC spectra of the isolated δ2 tail showed chemical shift perturbations upon addition of kinase domain alone (Fig. S2B, left) that were enhanced by the addition of ATP (Fig. S2B, right), suggesting that the δ2 tail interacts with the kinase domain in *trans* in a manner different from the δ1 tail. Peaks in the CON spectrum of the ^13^C,^15^N δ2 tail were significantly broadened compared to the δ1 tail (Fig. S2C, D), limiting further NMR analyses of the δ2 tail.

To determine how regulation of the δ1 tail is influenced when tethered to the kinase domain, we produced a segmentally isotopically labeled full-length kinase using an unlabeled kinase domain attached to an ^15^N-labeled δ1 tail through Sortase A-mediated ligation (Fig. S2E) (34, 35). ^15^N-^1^H HSQC spectra of the isolated ^15^N-labeled δ1 tail and the tethered tail appeared to be the same, aside from broadening of a single glycine (G296) near the ligation juncture (Fig. 2F). Importantly, this suggests that the kinase domain and tail do not interact in the unphosphorylated state in *cis*. Addition of ATP to the tethered tail sample resulted in chemical shifts in the ^15^N-labeled δ1 tethered tail backbone similar to those we observed in *trans* (compare Fig 2A and Fig. S2F), indicating that tethering did not substantially alter kinase domain interaction and phosphorylation of the tail, including activity on the XCT. To confirm this, we generated a minimal δ1 tail construct beginning at residue 376. We obtained chemical shift assignments for the phosphopeaks that correspond to the three phosphosites specific to the δ1 tail (Fig. 2G). Phosphorylation of these residues in the full-length CK1δ1 protein was confirmed by LC/MS-MS (Fig. S2G). Using ^15^N-edited 1D ^1^H spectra of the phosphoserine region (Fig. 2H), we were able to monitor the kinetics of phosphorylation of the δ1-specific residue S406. As expected, tethering increased the rate of phosphorylation compared to in *trans* (Fig. 2I) (36). Collectively, our NMR studies revealed differences in how the δ1 and δ2 tails interact with the kinase domain and demonstrated a preference for phosphorylation of the δ1-specific XCT serines by the kinase domain.

### Impact of phosphorylated FASP and CK1δ tail on structural dynamics of the kinase

Kinase domains interact with a number of ligands, from nucleotides and substrates to products and inhibitors, all of which can exert changes in dynamics or structure across the protein (37). To better understand autoinhibitory interactions between the kinase domain and tails of different CK1 isoforms, we turned to hydrogen-deuterium exchange mass spectrometry (HDX-MS) to characterize the conformational dynamics of different CK1δ isoforms. HDX-MS has been a powerful tool to provide insights into conformational dynamics in protein kinases (38–40). We first explored the effects of ATP or its non-hydrolyzable analogue AMPPNP binding on the isolated kinase domain using HDX-MS. Protection against deuterium incorporation upon binding to either nucleotide was predominantly observed across the N-terminal lobe and active site (Fig. S3A, D), with lower deuterium exchange in the bound state reflecting decreased solvent accessibility and increased H-bond formation between the nucleotide-interacting residues. At the active site, ATP binding led to stronger protection than AMPPNP (Fig. S3B). These effects are similar to those observed for other kinases including PKA, ERK2, IRK, and SRC (41–44). In addition to these direct effects, the binding of ATP and AMPPNP also elicited distal conformational changes across the C-terminal lobe. Reduced deuterium exchange was observed across peptides spanning the DFG motif (residues 145-151), the activation loop (151–179), and loop L-EF (208–218), which are all centered around the substrate-binding cleft/groove (Fig S3A). No significant protection was detected in other regions of the C-lobe. Comparison of HDX-MS results also revealed other intrinsic dynamics of CK1δΔC, wherein peptides spanning the catalytic cleft and hinge region (residues 56-72) displayed bimodal isotopic distribution of mass spectra (Fig. S3C). The two populations indicate the open-and closed-conformations of CK1δΔC that may be important for catalytic activity.

To validate HDX-MS as an approach to monitor inhibitory interactions on the kinase domain, we also looked at CK1δΔC in the presence of 4p-FASP, a phosphorylated inhibitory peptide from PER2 (20). Comparing HDX data of CK1δΔC in the absence and presence of 4p-FASP revealed a reduction in conformational flexibility throughout the kinase domain (Fig. 3A), with the largest effects observed at peptides covering the nucleotide-binding site (4–20), hinge region (56–72), activation loop (150–179), and substrate-binding cleft (residues 209-230) (Fig. 3B), close to two conserved anion binding sites that mediate a direct interaction with the 4p-FASP peptide (20). Comparing the effects of 4p-FASP binding to the HDX profile of ATP-bound CK1δΔC (Fig. 3A, B), we found that a larger segment of the activation loop and C-lobe had reduced solvent accessibility with 4p-FASP. These results illustrate how a phosphorylated product interacts with the substrate binding domain to substantially impact the conformational flexibility of CK1δ.

**Figure 3.**
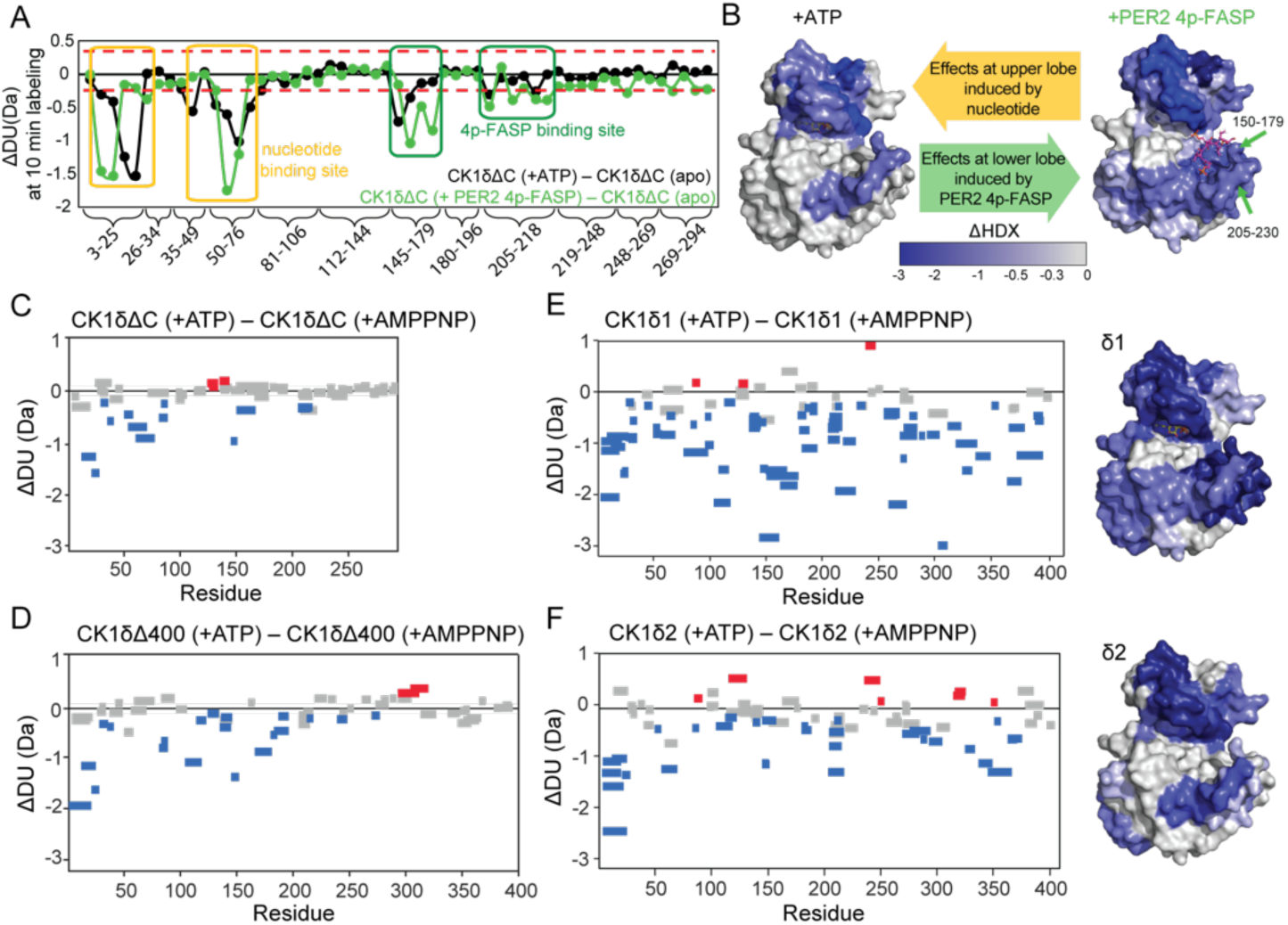
Impact of phosphorylated FASP and CK1δ tail on structural dynamics of the kinase. **A**, Comparison of differences in deuterium exchange of CK1δΔC in the presence and absence of 4p-FASP (green) or ATP (black) after 10 min labeling with residue numbers of the peptides grouped and indicated on the x-axis. Dashed line, significance threshold of ±0.28 Da. The deuterium exchange values are tabulated in Supplementary Table 1. **B**, Structural representation of deuterium exchange at 10 min with ATP (left, PDB: 6PXO, with ATP (yellow) from PDB: 6RU6) or 4p-FASP (right, PDB: 8D7O). **C-F**, Woods differential plots showing deuterium exchange between ATP-bound and AMPPNP-bound states of different CK1δ constructs: (**C**) CK1δΔC; (**D**) CK1δΔ400; (**E**) CK1δ1; and (**F**) CK1δ2 at 10 min labeling time. Data available in Supplementary Table 2. Peptides showing significant protection (blue), deprotection (red), or non-significant (gray) deuterium exchange based on ±0.28 Da cutoff (confidence interval 99.0%). **E-F,** Corresponding structural representation of deuterium exchange differences at 10 min of CK1δ1 and CK1δ2.

To explore how differences in isoform phosphorylation affect interactions within CK1δ, we preincubated CK1δΔC, CK1δΔ400, CK1δ1, and CK1δ2 for 15 min at 25°C with excess AMPPNP or ATP to block or promote autophosphorylation, respectively, and then compared deuterium exchange profiles by HDX-MS (Fig. 3C-F). The HDX-MS results, shown as Woods differential plots, highlight peptides that undergo significant protection from deuterium exchange in the presence of ATP in blue, while peptides with increased exchange are in red (Fig. 3C-F, S3E-H). Similar to nucleotide alone, we observed protection in CK1δΔC in peptides covering primarily the N-lobe and substrate-binding groove (Fig. 3C, S3E). Similar changes in HDX profiles were observed for CK1δΔ400, which lacks the XCT (Fig. 3D, S3F). Two peptides at the junction between the kinase domain and the tail (residues 293-312 and 304-320) experienced a small but significant increase in deuterium exchange, indicating increased flexibility of this region upon ATP binding, although no other significant changes were observed along the C-terminal tail of CK1δΔ400 (Fig. 3D S3F).

The presence of CK1δ1 and CK1δ2 XCTs led to different degrees of protection upon autophosphorylation (Fig 3E, F, S3G, H). In the kinase domain, CK1δ2 differed from CK1δΔ400 by modestly enhancing protection of the nucleotide binding pocket (Fig 3F S3H). However, CK1δ1, with its preferential phosphorylation sites in the XCT, was strikingly different. CK1δ1 autophosphorylation led to a large majority of peptides (71 peptides in CK1δ1 versus only 32 in CK1δ2) showing significant protection spanning the N-lobe and the C-lobe. Peptides spanning the length of the disordered tail on CK1δ1 also showed significant protection, suggesting solvent inaccessibility, possibly due to interdomain interactions. CK1δ2 also had many peptides in the C-terminal tail that showed protection against deuterium exchange, albeit to a lesser extent (Fig. 3E, F S3G,H).

To further understand changes in structural dynamics mediated by phosphorylation, we analyzed residue-specific deuterium exchange by analysis of overlapping peptides, represented as heat maps for each construct (Fig. S3I). These provide a comparative overview of time-dependent changes in exchange kinetics across the protein, including the C-terminal tail, and how truncation or splice isoforms influence this. These changes likely reflect both orthosteric effects of nucleotide-binding, as well as allosteric effects induced by the nucleotide and phosphorylation of the C-terminal tail. Lower overall deuterium exchange across CK1δ2 suggests that the δ2 tail has different conformational dynamics and/or interactions with the kinase domain from the δ1 tail, consistent with our NMR data. Furthermore, the effect of different tail lengths on the conformational changes on the kinase domain are clear, with increases in deuterium exchange observed across most peptides of the kinase domain as the CK1δ1 XCT was removed (Fig. S3J-M).

### Anion binding sites on the kinase domain play a role in autoinhibition by the δ1 tail

One defining feature of CK1 is anion binding sites near the active site that participate in substrate positioning and the conformation of the activation loop (22, 32). The phosphorylated PER2 pFASP peptide docks its phosphate groups into these sites that span the substrate binding cleft of the kinase, thus inhibiting kinase activity (Fig 4A, C) (20). We sought to determine if the phosphorylated XCTs utilize these anion binding sites to inhibit the kinase in a similar manner. First, we tracked changes in deuterium exchange of different kinase constructs in the presence and absence of ATP, looking specifically at several peptides that are proximal to anion binding site 1 (Fig. 4B) and site 2 (Fig. 4D). For anion binding site 1, CK1δ1 showed the greatest protection for all three peptides, with the 4pFASP-bound kinase, CK1δ2, and CK1δΔ400 showing moderate protection compared to CK1δΔC (Fig. 4B). For anion binding site 2, CK1δ1 and the 4pFASP-bound kinase showed greater protection than the others (Fig. 4D).

**Figure 4.**
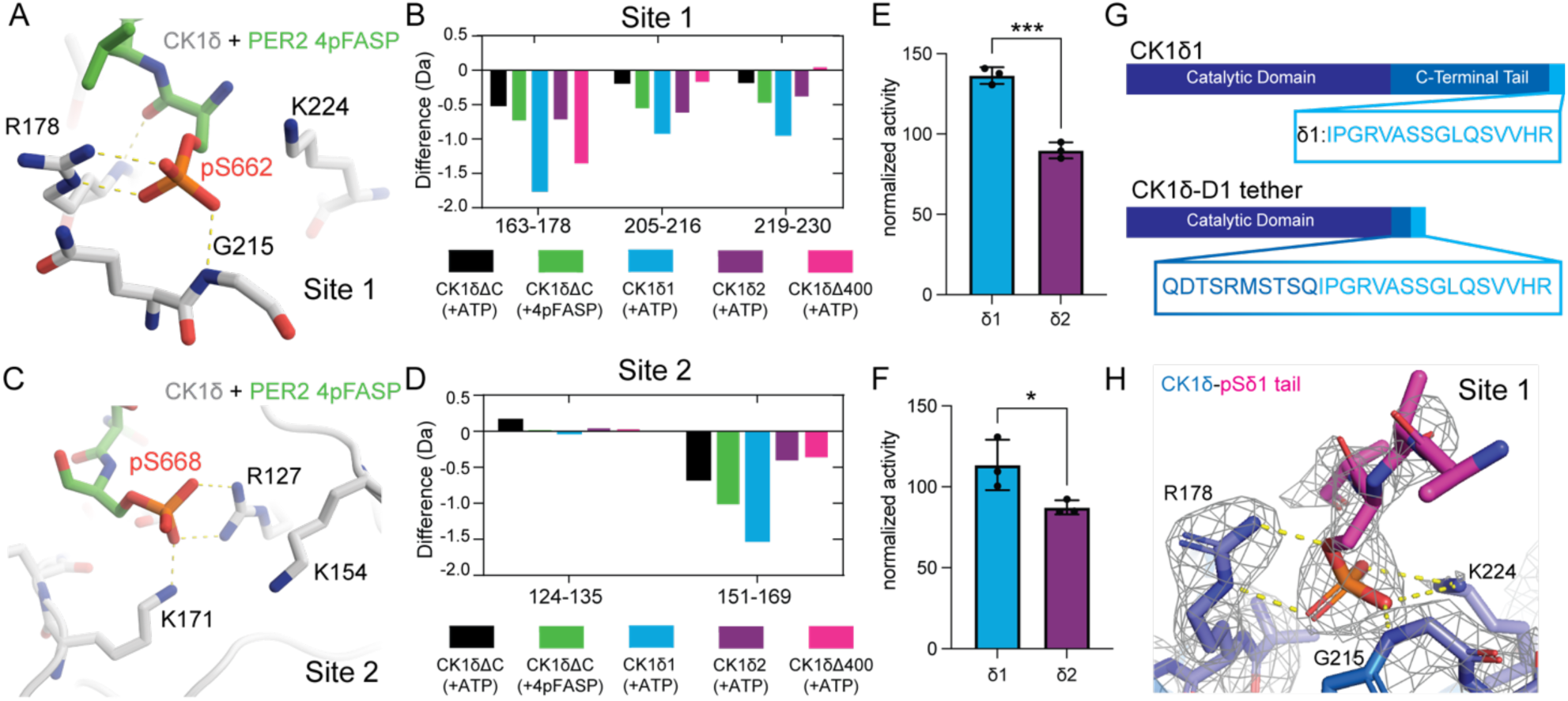
Anion binding sites on the kinase are required for autoinhibition by the δ1 tail. **A**, Structural view of CK1δ bound to the inhibitory PER2 4p-FASP (PDB: 8D70) showing phosphate coordination by anion binding site 1. **B**, Comparison of protection from deuterium exchange of peptides spanning anion binding site 1 with CK1δΔC (black), CK1δΔ400 (pink), CK1δ1 (blue), CK1δ2 (purple), and CK1δ-ΔC:4pFASP (green). **C**, Structural view of CK1δ bound to the inhibitory PER2 4p-FASP (PDB: 8D70) showing phosphate coordination by anion binding site 2. **D**, Comparison of protection from deuterium exchange of peptides spanning anion binding site 2. **E-F**, ^32^P kinase assays comparing the activity of K224D (**E**) or R127E (**F**) in CK1δ1 (blue) and CK1δ2 (purple) to WT, normalized to the effect of the mutant on the isolated kinase domain (Fig. S5) (n=3, mean ± s.d.). Significance assessed by unpaired two-tailed t-test: *, p<0.05; ***, p<0.001. **G,** Schematic of full length CK1δ1 and CK1δ-D1 tether construct used for crystallization of the kinase with phosphorylated XCT. **H,** 2Fo-Fc omit map of anion binding site 1 (PDB: 9B3S) contoured at 1 σ highlighting residues on the kinase domain (blue), the phospho-tail (magenta), and polar contacts (yellow dashed lines).

To determine if these sites play a role in inhibition by the tails, we looked at the effect of a single, well-characterized point mutation at each anion binding site (22) on the ability of CK1δ1 or CK1δ2 to phosphorylate the PER2 FASP priming serine. Notably, these anion binding site mutations eliminated inhibition by the phosphorylated PER2 FASP region (20). The K224D mutation impaired phosphorylation by CK1δ2 and CK1δΔC, which we attribute to impaired substrate positioning based on prior work (22). Remarkably, the same mutant in CK1δ1 had no decrease in FASP priming (Fig. 4E, S4A-B). Likewise, disruption of site 2 (R127E) had a similar isoform-specific effect on autoinhibition (Fig. 4F, S4C-D), although the effect was more modest than for site 1. We propose that the impaired substrate positioning caused by mutation of the anion binding sites is counter-balanced by relief of autoinhibition in CK1δ1, perhaps due to reduced XCT binding at the mutated anion binding sites.

To show direct interaction of the phosphorylated XCT with the anion binding sites, we generated a CK1δ1 construct with a minimal tail attached to the kinase domain that we called CK1δ-D1 tether (Fig. 4G). Kinase assays comparing CK1δ1, CK1δΔC, and CK1δ-D1 tether on PER2 FASP S665A show the tethered tail retains moderate inhibition (Fig. S4E, F). We then solved a crystal structure of the autophosphorylated CK1δ-D1 tether, observing density for a phosphorylated serine docked into anion site 1. There was poor density for the sidechains of surrounding residues on the peptide, impeding the identification of the exact phosphor-serine(s) bound (Fig. 4H). This shows that the XCT directly interacts with anion site 1 in a similar manner to the inhibitory 4p-FASP (Fig. 4A, H), demonstrating a conserved mechanism of inhibition.

### Isoform-specific sequences of CK1δ differentially inhibit the kinase domain

Prior studies have shown that at least two substrates phosphorylated by CK1, the PER2 FASP region and the phosphorylation activation domain (PAD) of p63, inhibit kinase activity by product inhibition (20, 21). To test if this model holds for CK1δ tail autoinhibition, we added different phosphorylated δ1 or δ2 peptides to the kinase in *trans* (Fig. S5B). To look at their effect on kinase activity, we used ADP-Glo assays with two PER2 substrates, the FASP and the Degron, measuring activity with increasing phosphopeptide. For both substrates, inhibition by the δ1 peptides relied more on phosphorylation of the XCT, and the phosphorylated δ1 peptides were generally more potent inhibitors than the δ2 peptides, although there were some minor differences between the two substrates (Fig. 5A, S5A).

**Figure 5.**
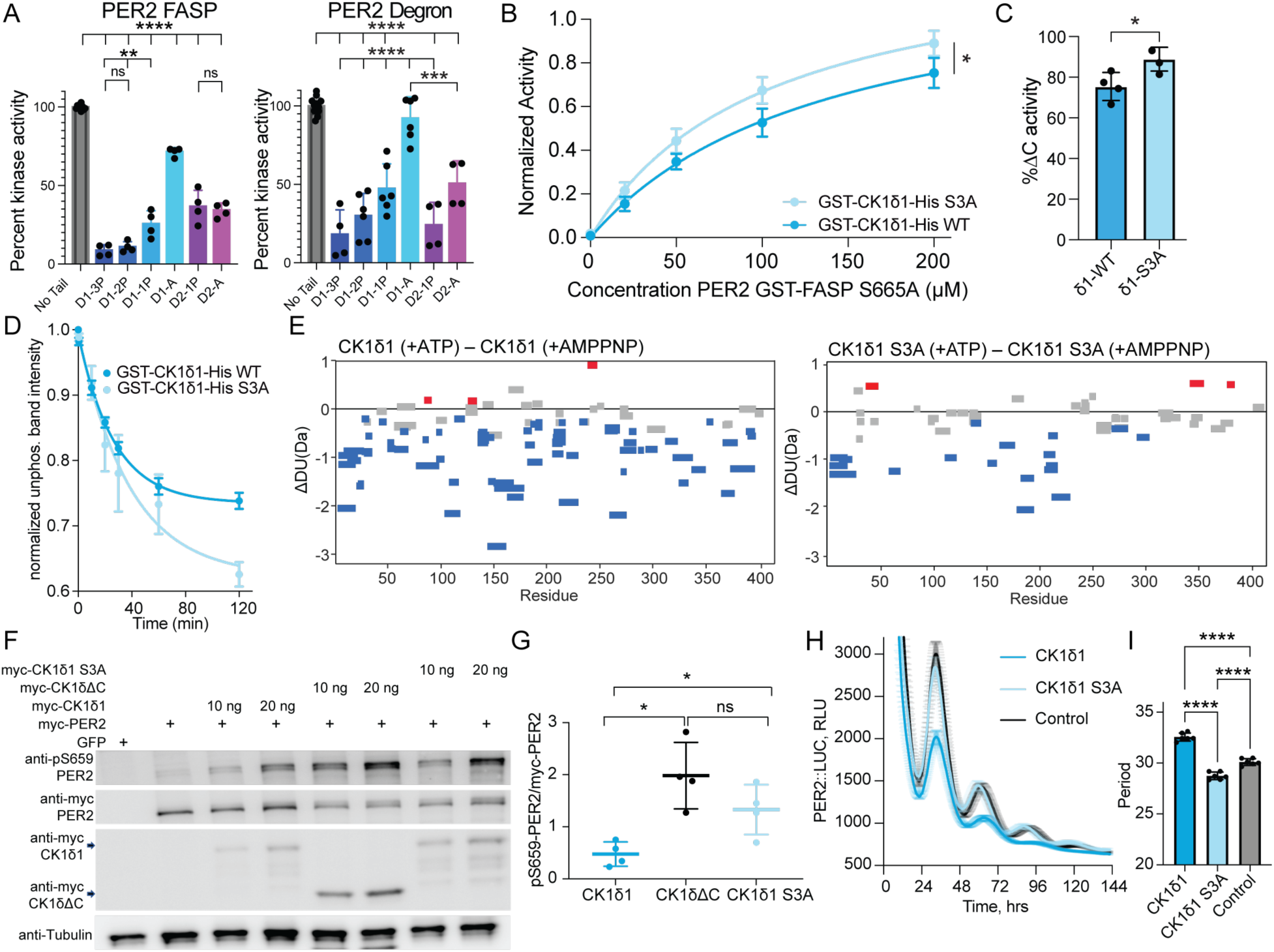
Variable C-terminal tail length and mutations impacts kinase activity of CK1δ on its substrate PER2. **A**, ADP-Glo kinase assay with phosphorylated CK1δ C-terminal peptides on PER2 FASP (left) or Degron (right). Significance assessed by ordinary one-way ANOVA **, p<0.01; ****, p<0.0001 (n=2, mean ± s.d.). **B**, ^32^ P kinase assay of CK1δ1 (blue) or the S3A mutant (light blue) on PER2 GST-FASP-S665A. Significance assessed by Extra sum-of-squares F test (n=3 mean ± s.d.). **C**, Quantification of the endpoint (200μM) in panel C (n=3 mean + s.d.) Significance assessed by unpaired two-tailed t-test: *, p<0.05. **D,** Quantification of the unphosphorylated band intensity over time for CK1δ1 (blue) and CK1δ1 S3A (light blue) normalized to time 0 band intensity (n=3, mean +/-SEM). **E**, Woods differential plots showing deuterium exchange between ATP-bound and AMPPNP-bound states of CK1δ1 (top) and CK1δ1 S3A (bottom) at 10 min labeling. Data available in Supplementary Table 2. Peptides showing significant protection (blue), deprotection (red) or non-significant (gray) deuterium exchange based on ±0.28 Da cutoff (confidence interval 99.0%). **F**, Representative western blot of mouse PER2 FASP priming phosphorylation (pS659) in HEK 293 lysates expressing indicated proteins (n=4). **G**, The ratio of pS659 to myc-PER2 was calculated and values are expressed in y-axis. (n=4 with mean ± s.d.). Significance assessed by unpaired two-tailed t-test: *, p<0.05. **H,** Luminescence trace (shown in RLU) of PER2::LUC U2OS cells with stably expressed CK1δ1 (blue), CK1δ1 S3A (light blue), or no kinase control (black). (n=6, +/-SEM). Representative trace of n=3. **I,** Period analysis of panel I showing a lengthened period for CK1δ1 (blue) and a shortened period for CK1δ1 S3A (light blue) compared to control cells (black). Significance assessed by ordinary one-way ANOVA, ****, p<0.0001. (n=6, mean +/-s.d.).

We extended these findings to full-length CK1δ1 and tested if the δ1-specific phosphosites are necessary for inhibition in the full-length protein *in vitro*. We mutated the three unique phosphosites in δ1 to alanines (S3A: S406A/S407A/S411A) and compared the activity to wild-type CK1δ1 on the PER2 FASP S665A substrate (Fig. 5B, S5C). Consistent with the D1-A peptide data, the CK1δ1 S3A mutant showed a modest but significant increase in activity compared to CK1δ1 (Fig. 5B, C), demonstrating that phosphorylation of the extreme C-terminus is important for autoinhibition of CK1δ1. Coomassie gel shift assays also showed that CK1δ1 S3A shows a higher rate of autophosphorylation (Fig. 5D, S5D), indicating the XCT also inhibits upstream autophosphorylation of other sites on the tail.

We next tested if the S3A mutant altered the interaction and dynamics of the tail with the kinase domain as assessed by HDX-MS. Strikingly, we found a large decrease in protected peptides in CK1δ1 S3A relative to the wild-type enzyme (from 71 to 19, Fig. 5E, S5E), spanning both the kinase domain and the C-terminal tail. The protection of the kinase domain by CK1δ1 S3A most closely resembled the HDX-MS behavior for CK1δΔC (Fig. 3C), suggesting a near-total loss of tail-kinase interaction in the mutant. Hence, phosphorylation of three serines in the CK1δ1 XCT is likely to be a prerequisite for the interaction of the C-terminal tail with the kinase domain.

These findings led us to examine how the CK1δ1 S3A mutation affected PER2 FASP phosphorylation and circadian rhythms in cells. Deletion of the C-terminal tail of CK1δ1 (CK1δΔC) or its S3A mutation markedly increased FASP priming site phosphorylation in full-length PER2 in cells (Fig. 5F, G). To test how the S3A mutations influence circadian rhythms, we created PER2::LUC U2OS cell lines with stable overexpression of Flag-tagged CK1δ1 or the S3A mutant (Fig. S5F). Expression of CK1δ1 S3A shortened period by ∼3 hrs relative to cells expressing CK1δ1 WT (Fig. 5H, I), consistent with its overall increase in kinase activity. These results demonstrate that phosphorylation of the three CK1δ1-specific serines in the XCT is important for regulation of kinase activity in cells, where it contributes to period determination.

## Discussion

CK1δ has been considered a constitutively active kinase because it does not require activation loop phosphorylation to take on an active conformation. Instead, it relies on autophosphorylation of its C-terminal tail and/or feedback inhibition from phosphorylation of its substrates to regulate kinase activity (3, 4, 20). While we are beginning to understand mechanisms of feedback inhibition (20, 21), the extent of intrinsic disorder and phosphorylation throughout the tail have impeded a mechanistic understanding of autoinhibition. Changes in circadian rhythms with two isoforms of CK1δ that differ only in the extreme C-terminus of the tail demonstrate unambiguously that the tails have an important role *in vivo* (26, 27). However, it was not clear from these studies if isoform-specific differences in activity were due to regulation of the tail by other kinases or if they were intrinsic to CK1δ. Here, we show that CK1δ isoforms have inherent differences in activity arising from interactions of the kinase domain and its disordered tail that are specified by isoform-specific differences in autophosphorylation of the XCT. These differences explain the changes in period seen in the different CK1δ splice variants.

NMR spectroscopy allowed us to track site-specific autophosphorylation of the CK1δ disordered tails. We found that the 3 serines of the δ1-specific XCT are preferentially phosphorylated over other sites in the tail, whether the tail is phosphorylated by the kinase in *trans* or covalently linked to the kinase domain in *cis*. Chemical shift perturbations provided strong evidence for phosphorylation-independent effects of the kinase domain on the δ2 tail, although further analysis of the δ2 tail by NMR was limited due to extensive peak broadening in the absence of the kinase domain. This behavior is consistent with potential “fuzzy” intramolecular interactions within the δ2 tail, which may exhibit multiple transient conformations, as has been shown for other intrinsically disordered proteins (45, 46). Isoform-specific differences in the dynamics of the disordered tails could contribute to the decreased autoinhibition we observed with CK1δ2. Due to the sequence similarity of the XCT between CK1δ2 and CK1ε, the differences in tail dynamics we observed here could extend to CK1ε. Furthermore, the presence of disordered N-or C-terminal tails in CK1δ orthologs of other species also suggests that this could be a conserved mechanism of autoinhibition.

The phosphorylated PER2 FASP substrate inhibits CK1δ through interaction with conserved anion binding sites that flank the substrate binding cleft to regulate circadian rhythms (20). HDX-MS analysis of the kinase domain bound to the inhibitory 4p-FASP phosphopeptide showed increased protection from deuterium exchange near these sites, confirming that our HDX-MS pipeline could map inhibitory interactions on the kinase domain. Of the two CK1δ isoforms, CK1δ1 showed the greatest deuterium protection across the kinase domain, including at sites protected by the 4p-FASP peptide. Protection in the kinase domain was moderately reduced in CK1δ2 and essentially eliminated by truncation of the XCT or the S3A mutant that eliminated phosphosites unique to δ1, indicating that phosphorylation of the XCT plays a crucial role in autoinhibition by CK1δ1. We also observed significant protection of the intrinsically disordered tail upon autophosphorylation of both CK1δ1 and CK1δ2 that was eliminated with truncation of the isoform-specific sequences. This suggests that both tails make extensive contact with the kinase domain, although we don’t yet understand the specific details of these interactions. One recent study looked at autoinhibitory interactions of CK1δ1 in the context of kinase activity on p63, finding no requirement for any specific phosphorylation sites (47), while another demonstrated that different modes of autoinhibition by CK1δ and ε isoforms influences substrate specificity (30), perhaps by targeting different sites on the kinase. Although our crystal structure of the tethered δ1 XCT revealed a key role for site 1 in kinase binding, we were unable to resolve which phosphor-serine(s) of the δ1 XCT were bound.

Based on our results, we propose that autophosphorylation of the CK1δ tail in its XCT facilitates its interaction with the kinase domain to inhibit kinase activity. Here, we examined the effect of CK1δ autophosphorylation on PER2 substrates involved in a phosphoswitch that controls its stability and the period of circadian rhythms in mammals, including humans (12, 13). The phosphoswitch alone does not fully explain the difference in period displayed by over-expression of CK1δ1 WT and the S3A mutant; however, recent reports identified a role for CRY-PER-CK1δ complexes in phosphorylating the CLOCK subunit of the circadian transcription factor CLOCK:BMAL1 to suppress transcriptional activation of target genes by displacing the complex from DNA (15, 48, 49). Therefore, regulation of CK1δ activity by isoform-specific tails could have several roles in regulating circadian rhythmicity, not only through modulating the stability of PER2 but also by instigating transcriptional repression through promoting the release of CLOCK:BMAL1 from DNA. CK1 isoforms also have major roles in other pathways, such as apoptotic signaling (50, 51), Wnt signaling (52–54), and the cell cycle, where it is overexpressed in several cancers (55, 56). Interestingly, CK1 autophosphorylation is stabilized in a cell cycle-dependent manner (57), suggesting that understanding its regulatory mechanisms could begin to build a deeper understanding of this enigmatic kinase.

## STAR Methods

### RESOURCE AVAILABILITY

#### Lead contact

Further information and requests for resources and reagents should be directed to and will be fulfilled by the lead contact, Carrie L. Partch.

#### Materials availability

- Plasmids generated in this study are available upon request.

#### Data availability

- Data generated in this study is included in the manuscript.
- NMR chemical shift assignments for the human CK1 δ1 tail (accession # 52349) and δ2 tail (accession #52375) have been deposited at BMRB.
- Coordinates for the CK1δ1-D1 tether structure can be found at the PDB with accession # 9B3S.

### EXPERIMENTAL MODEL AND SUBJECT DETAILS

#### Cell lines

- The HEK293T cell line was purchased from ATCC (#CRL-3216).
- U2OS PER2::LUC cell line was a gift from John O’Neill (MRC LMB, UK).

## METHOD DETAILS

### Protein constructs, expression, and purification

All constructs were expressed from a pET22b vector backbone containing TEV-cleavable tags generated in-house based off the Parallel vector series (58), and have an N-terminal vector artifact (GAMDPEF) remaining after TEV cleavage. Mutations were introduced using the site-directed mutagenesis protocol in (59) and validated by sequencing. The CK1δ-D1 tether sequence was synthesized (Twist Biosciences) and inserted into in-house vectors using standard cloning.

Full-length human CK1δ isoforms (CK1δ1, residues 1-415; CK1δ2, residues 1-409) and CK1δ Δ400 (residues 1-400) were expressed with an N-terminal TEV-cleavable GST tag and a C-terminal His tag. Different constructs of the CK1δ catalytic domain, CK1δ ΔC (residues 1-294 or 1-317) and CK1δ-D1 tether were expressed with an N-terminal TEV-cleavable His-GST tag. A construct of the kinase domain for Sortase A-mediated ligations, CK1δ ΔC*, containing residues 1-294 with a C-terminal Sortase A recognition site (LPETGG) was expressed with an N-terminal TEV-cleavable His-MBP tag. Constructs of isolated CK1 tails (CK1δ1 tail, residues 295-415; CK1δ2 tail, residues 295-409; CK1δ1 short tail, residues 376-415) were expressed with an N-terminal TEV-cleavable His-NusA tag (HNXL) with an N-terminal tryptophan added for UV detection during purification.

Recombinant human PER2 peptides (FASP peptide, residues 645–687; Degron peptide, residues 475–505) were expressed with an N-terminal TEV-cleavable His-NusA tag when used for the ADP-Glo peptide inhibition assays or with an N-terminal TEV-cleavable His-GST tag for the radioactive kinase assay. All recombinant PER2 peptides contain an N-terminal tryptophan for UV detection during purification and a polybasic motif (WRKKK) following the vector artifact, as in (20).

GST-lambda (λ) phosphatase-Strep-tag II was expressed with a TEV-cleavable N-terminal His-GST tag and a C-terminal Strep-tag II. His-Sortase A 7M (Addgene # 51141) and GST-λ phosphatase were expressed in BL21 (DE3) Escherichia coli, while all CK1 and PER constructs were expressed in Rosetta2 (DE3) E. coli. In general, cells were grown in LB media (for natural abundance growths) or M9 minimal media with the appropriate stable isotopes (i.e., 15N/13C) for NMR as done before (27) at 37°C with shaking until the OD600 reached ∼0.6-0.8. Expression was induced with 0.5 M IPTG, and cells were grown for an additional 16-20 hours at 18°C.

For purification of full-length CK1δ, cells were lysed in 50 mM Tris pH 7.5, 500 mM NaCl, 5% glycerol, 1 mM EDTA, 1 mM TCEP, 0.05% Tween using a high-pressure homogenizer. GST-tagged full-length proteins were purified from soluble lysate using Glutathione Sepharose resin (Cytiva) and eluted with buffer containing 50 mM Tris pH 7.5, 500 mM NaCl, 5% glycerol, 1 mM EDTA, 1 mM TCEP, 0.05% Tween, and 25 mM reduced glutathione. To remove phosphorylation from the kinase that may have occurred during expression or purification, GST-tagged purified proteins were treated with 200 nM GST-λ phosphatase-Strep tag II supplemented with 10 µM MnCl_2_ in λ phosphatase buffer (50 mM HEPES pH 7.5, 100 mM NaCl, 0.1 mM EDTA, 1 mM TCEP) overnight at 4°C. Strep-Tactin Sepharose resin (Cytiva) was used to remove GST-λ phosphatase-Strep tag II, and dephosphorylated CK1δ was collected with several washes of 50 mM Tris pH 7.5,

500 mM NaCl, 5% glycerol, 1 mM EDTA, 1 mM TCEP, and 0.05% Tween. Dephosphorylated proteins were then further purified by the C-terminal His-tag using Ni-NTA resin (Qiagen). Full-length dephosphorylated proteins were subsequently purified by size-exclusion chromatography on a HiLoad 16/600 Superdex 200 prep grade column (Cytiva) in 50 mM Tris pH 7.5, 200 mM NaCl, 1 mM EDTA, 5 mM β-mercaptoethanol (BME), 5% glycerol, and 0.05% Tween, and then frozen in aliquots for storage at -70°C.

For CK1δ catalytic domain preps, cells were lysed in 50 mM Tris pH 7.5, 500 mM NaCl, 5% glycerol, 1 mM EDTA, 1 mM TCEP, 0.05% Tween using a high-pressure homogenizer. His-GST-tagged proteins were purified from soluble lysate using Glutathione Sepharose resin (Cytiva) and eluted with 50 mM Tris pH 7.5, 500 mM NaCl, 5% glycerol, 1 mM EDTA, 1 mM TCEP, 0.05% Tween containing 25 mM reduced glutathione. To cleave the His-GST tag, His-TEV protease was incubated with the protein at 4°C overnight. Cleaved CK1δ ΔC was purified away from His-GST and His-TEV using

Ni-NTA resin (Qiagen) and size-exclusion chromatography on a HiLoad 16/600 Superdex 75 prep grade column (Cytiva) in 25 mM MOPS pH 7.0, 1 mM EDTA, 100 mM NaCl, 5 mM BME, 11 mM MgCl_2_, and 0.05% Tween, and then frozen in aliquots for storage at -70°C.

For CK1δ-D1 tether preps, cells were lysed in 50 mM Tris pH 7.5, 500 mM NaCl, 5% glycerol, 1 mM EDTA, 1 mM TCEP, 0.05% Tween using a high-pressure homogenizer. His-GST-tagged proteins were purified from soluble lysate using Glutathione Sepharose resin (Cytiva) and eluted with 50 mM Tris pH 7.5, 500 mM NaCl, 5% glycerol, 1 mM EDTA, 1 mM TCEP, 0.05% Tween containing 25 mM reduced glutathione. To cleave the His-GST tag, glutathione was diluted out and GST-TEV protease was incubated with the protein at 4°C overnight. Cleaved CK1δ-D1 tether was purified away from His-GST and GST-TEV using Glutathione Sepharose resin (Cytiva). Protein was collected from the flow-through, then further purified using ion exchange chromatography with a HiTrap SP XL column (Cytiva). Protein for crystal trays was then given 1 mM ATP and 10mM MgCl_2_ to autophosphorylate at 4°C overnight. It was then further purified with size-exclusion chromatography on a HiLoad 16/600 Superdex 75 prep grade column (Cytiva) in 25 mM MOPS pH 7.0, 1 mM EDTA, 100 mM NaCl, 5 mM BME, 11mM MgCl_2_, and 0.05% Tween, and then frozen in aliquots for storage at -70°C.

For CK1δ ΔC* preps, cells were lysed in 50 mM Tris pH 7.5, 500 mM NaCl, 2 mM TCEP, 5% glycerol and 25 mM imidazole using a high-pressure extruder homogenizer. His-MBP-CK1δ ΔC* was purified using Ni-NTA resin (Qiagen) using standard approaches and eluted from the resin using 50 mM Tris pH 7.5, 500 mM NaCl, 2 mM TCEP, 5% glycerol and 250 mM imidazole. His-TEV protease was added to cleave the His-MBP tag from the CK1δ ΔC* at 4°C overnight. The cleavage reaction was subsequently concentrated and diluted to low imidazole concentration (< 25 mM), and then the kinase was purified away from His-MBP and His-TEV using Ni-NTA resin with 50 mM Tris pH 7.5, 500 mM NaCl, 2 mM TCEP, 5% glycerol and 25 mM imidazole. Protein was collected from the flow-through, then further purified using ion exchange chromatography with a HiTrap SP XL column (Cytiva) preceding size-exclusion chromatography on a HiLoad 16/600 Superdex 75 prep grade column (Cytiva) in 25 mM MOPS pH 7.0, 1 mM EDTA, 100 mM NaCl, 5 mM BME, 11mM MgCl_2_, and 0.05% Tween, and then frozen in aliquots for storage at -70°C.

For CK1δ1 tail and CK1δ2 tail preps, cells were lysed in 50 mM Tris pH 7.5, 500 mM NaCl, 2 mM TCEP, 5% glycerol and 25 mM imidazole using a high-pressure extruder homogenizer. His-NusA-tail fusion proteins were purified using Ni-NTA resin (Qiagen) using standard approaches and His-TEV protease was added to cleave the His-NusA tag from the tail on column at 4°C overnight. Tails were purified away from the resin containing His-NusA and His-TEV with 50 mM Tris pH 7.5, 500 mM NaCl, 2 mM TCEP, 5% glycerol and 25 mM imidazole. Tails were further purified by size exclusion chromatography on a HiLoad 16/600 Superdex 75 prep grade column (Cytiva) using NMR buffer (25 mM MOPS pH 7.0, 1 mM EDTA, 100 mM NaCl, 5 mM BME, 11 mM MgCl_2_, and 0.05% Tween) and frozen in aliquots for storage at -70°C.

For PER2 peptide preps, cells were lysed in 50 mM Tris pH 7.5, 500 mM NaCl, 2 mM TCEP, 5% glycerol and 25 mM imidazole using a high-pressure extruder homogenizer. His-NusA-FASP or His-NusA-Degron fusion proteins were purified using Ni-NTA resin (Qiagen) using standard approaches and His-TEV protease was added to cleave the His-NusA tag from the PER2 peptides on column at 4°C overnight. Peptides were purified away from the resin containing His-NusA and His-TEV with 50 mM Tris pH 7.5, 500 mM NaCl, 2 mM TCEP, 5% glycerol and 25 mM imidazole. Peptides were purified by size exclusion chromatography on a HiLoad 16/600 Superdex 75 prep grade column (Cytiva) using 1X kinase buffer (25 mM Tris pH 7.5, 100 mM NaCl, 10 mM MgCl_2_, and 2 mM TCEP). For His-GST-tagged PER2 peptides used in the radioactive kinase assays, cells were lysed in 50 mM Tris pH 7.5, 500 mM NaCl, 5% glycerol, 1 mM EDTA, 1 mM TCEP, 0.05% Tween using a high-pressure homogenizer. His-GST-tagged proteins were purified from soluble lysate using Glutathione Sepharose resin (Cytiva) and eluted with 50 mM Tris pH 7.5, 500 mM NaCl, 5% glycerol, 1 mM EDTA, 1 mM TCEP, 0.05% Tween containing 25 mM reduced glutathione. The peptides were further purified using size-exclusion chromatography on a HiLoad 16/600 Superdex 75 prep grade column (Cytiva) in 25 mM Tris pH 7.5, 100 mM NaCl, 10 mM MgCl_2_, and 2 mM TCEP, and then frozen in aliquots for storage at -70°C.

For His-Sortase A 7M, cells were lysed using a high-pressure extruder homogenizer followed by brief sonication at 4°C. After clarifying lysate on a centrifuge at 4°C at 140,500 × g for 1 hour, protein was captured using Ni-NTA affinity chromatography (Qiagen). The His-tag was cleaved using His-TEV protease on Ni-NTA resin at 4°C overnight. Cleaved protein was then collected from the flow-through and further purified using size exclusion chromatography on a HiLoad 16/600 Superdex 75 prep grade column (Cytiva) in 50 mM Tris, pH 7.5, 150 mM NaCl, and 10% glycerol, and then frozen in aliquots for storage at -70°C.

For GST-λ phosphatase-Strep tag II, cells were lysed in 50 mM Tris pH 7.5, 500 mM NaCl, 5% glycerol, 1 mM EDTA, 1 mM TCEP, 0.05% Tween using a high-pressure homogenizer. Proteins were purified from soluble lysate using Glutathione Sepharose resin (Cytiva) and eluted with a solution containing 50 mM Tris pH 7.5, 500 mM NaCl, 5% glycerol, 1 mM EDTA, 1 mM TCEP, 0.05% Tween, and 25 mM reduced glutathione. λ-phosphatase was further purified using size-exclusion chromatography on a HiLoad 16/600 Superdex 75 prep grade column (Cytiva) in 50 mM HEPES pH 7.5, 100 mM NaCl, 0.1 mM EDTA, and 1 mM TCEP, and then frozen in aliquots for storage at -70°C.

### Solid-phase synthesis of human PER2 and CK1δ peptides

The following peptides were synthesized via solid-phase methods and purified to >90% purity by Biopeptide, Co.: human PER2 4pFASP, GKAEpSVApSLTpSQCpSYA (where pS represents a phosphoserine); human CK1δ D1-1P, pGlu-IPGRVAAAGLQpSVVHR; human CK1δ D1-2P, pGlu-IPGRVApSpSGLQAVVHR; human CK1δ D1-3P, pGlu-IPGRVApSpSGLQpSVVHR; human CK1δ D1-A, pGlu-IPGRVAAAGLQAVVHR; human CK1δ D2-1P, pGlu-NpSIPFEHHGK; human CK1δ D2-A, pGlu-NAIPFEHHGK. Following peptides used for ELISA assay were synthesized by Genscript with ≥95% purity, human CK1-δ1 tail peptide: IPGRVASSGLQSVVHR, human CK1-δ1-3P-tail: IPGRVApSpSGLQpSVVHR.

### Segmental labeling with Sortase A

To generate the segmentally isotopically labeled kinase, we utilized Sortase A-mediated ligation reactions (35) between a natural abundance kinase domain (CK1δ ΔC*, harboring the Sortase A recognition motif, LPETGG) and a uniformly ^15^N-labeled CK1δ1 tail with a free N-terminal glycine left after TEV cleavage. The reaction was incubated and dialyzed overnight at 4°C in (50 mM Tris pH 7.5, 200 mM NaCl, 1 mM EDTA, 5 mM BME, and 0.05% Tween). The His-Sortase A 7M enzyme was purified away from the reaction using Ni-NTA resin (Qiagen) with 50 mM Tris pH 7.5, 500 mM NaCl, 2 mM TCEP, 5% glycerol and 25 mM imidazole, while the ligated CK1δ ΔC*-^15^N CK1δ1 tail product from the flow-through was further purified by size-exclusion chromatography on a Superdex 75 Increase 10/300 GL analytical grade column (Cytiva) in NMR buffer (25 mM MOPS pH 7.0, 1 mM EDTA, 100 mM NaCl, 5 mM BME, 11 mM MgCl_2_, and 0.05% Tween) and frozen in aliquots for storage at -70°C.

### *In vitro* kinase activity assays

#### ɣ^32^P-ATP kinase assay

Radioactive kinase assays were performed using human PER2 His-GST-FASP S665A or GST alone as the substrate. Reaction mixtures were prepared in reaction buffer (25 mM Tris pH 7.5, 200 mM NaCl, 10 mM MgCl_2_, and 1 mM DTT) with 1 µM enzyme and 0-200 µM dilutions of the substrate as indicated at 25°C. Reactions were initiated with the addition of 2 mM ATP containing 2 μCi of ɣ-32p ATP (Perkin Elmer). All reactions were quenched with 2X SDS buffer after an incubation time of 1 hour. SDS-PAGE was performed on samples for each reaction, and the gels were dried at 80°C for 2 hours before being transferred to a storage phosphor screen (Amersham Biosciences) overnight. Images were collected with Typhoon Trio (Amersham Biosciences) and data were analyzed by densitometry using Image J (NIH), Excel (Microsoft), and Prism (GraphPad).

#### ELISA-based kinase assay

2 μg/mL of FASP-WT peptide was diluted in carbonate buffer, pH 9.5 and coated onto a 96 well plate (100 μL/well). The next day, 50 ng of CK1δ1-FL (from Signalchem) was incubated in kinase buffer (25 mM Tris pH 7.5, 5 mM beta glycerol phosphate, 2 mM DTT and 0.1 mM sodium orthovanadate) with or without 10 mM magnesium chloride and 200 μM ATP to allow for autophosphorylation for 30 min at 25°C in a reaction mixture of 20 µL. ELISA plate wells were washed thrice with wash buffer (PBS with 0.05% Tween 20, PBS-T) and once with kinase buffer. Reactions containing CK1δ1-FL purified protein with or without autophosphorylation were added to each well and the plate was incubated at 30°C for 1 hour. Next, the reaction mixture was discarded, and wells were washed with wash buffer thrice and incubated with blocking buffer (PBS-T with 5% BSA) for 1 hour at room temperature. Subsequently, wells were incubated with the polyclonal pS659 Ab, an anti-rabbit antibody conjugated to Biotin, and Streptavidin-HRP for 1 hour each at room temperature with washing as above after each incubation. For signal detection, TMB reagent (1-Step Ultra TMB-ELISA, Thermo Scientific) was added, incubated at room temperature for color development and quenched by addition of STOP solution (Thermo Scientific). The plate was read at 450 nm using an xMark Spectrophotometer plate reader (BioRad). Each sample was analyzed in triplicate.

#### ADP-Glo peptide inhibition assay

Kinase reactions were performed on the recombinant human PER2 FASP or Degron using the ADP-Glo kinase assay kit (Promega) according to manufacturer’s instructions. Synthetic peptide inhibitors were solubilized in water at a concentration of 10 mM. All kinase reactions were performed in 30 μL reactions in duplicate (n = 2 independent assays) with 100 µM ATP, 0.2 µM CK1δ ΔC kinase, and 50 µM FASP or 100 µM Degron substrates were incubated in 1X kinase buffer (25 mM Tris pH 7.5, 100 mM NaCl, 2 mM TCEP, 10 mM MgCl_2_) at room temperature for 1 hour for FASP or 3 hours for Degron in the presence of increasing amounts of synthetic δ1 or δ2 peptides as indicated. 5 µL aliquots were quenched with ADP-Glo reagent after incubation, and luminescence measurements were taken at room temperature with a SYNERGY2 microplate reader (BioTek) in black opaque 384-well microplates. Data analysis was performed using Prism (GraphPad).

#### Coomassie Gel Shift

Gel shift kinase assays were performed using 1 µM of indicated CK1 isoform in 1X kinase buffer (25 mM Tris pH 7.5, 100 mM NaCl, 2 mM TCEP, 10 mM MgCl_2_). Reactions were initiated with addition of 2mM ATP and quenched at indicated time points with 2X SDS. SDS-PAGE was performed on samples for each reaction, and the gels were stained with SimplyBlue™ SafeStain (Invitrogen), destained overnight, and then imaged with ChemiDoc Imaging System (Bio-Rad). Analysis was performed with ImageJ, where the relative pixel density of each band was normalized to the corresponding band at 0 mins of the reaction.

### Crystallization and structure determination

Crystallization was performed by hanging-drop vapor-diffusion method at 22°C by mixing an equal volume of protein with reservoir solution. The reservoir solution for CK1δ-D1 tether (5.5 mg/mL) was 0.21 mM succinic acid pH 5.5 and 17% (vol/vol) PEG 3350. The crystals were looped and briefly soaked in a drop of cryopreservant reservoir solution with 20% (vol/vol) glycerol and then flash-frozen in liquid nitrogen for X-ray diffraction data collection. Data sets were collected at the 23-ID-D beamline at the Advanced Photon Source (APS) at the Argonne National Laboratory. Data were indexed, integrated, and merged using CCP4 software suite (60). Structures were determined by molecular replacement with Phaser MR (61) using anion free CK1δΔC (PDB: 6PXO). Model building was performed with Coot (62) and structure refinement was performed with PHENIX (63). All structural models were generated using PyMOL Molecular Graphics System 2.5.4 (Schrödinger).

### ELISA, Autophosphorylation, and Western Blotting

For ELISA, 1 µg/ml of indicated tail peptide was diluted in carbonate buffer, pH 9.5 and coated onto a 96 well ELISA plate (100 µl/well) and incubated overnight at 4. The next day, wells were washed with wash buffer (PBS with 0.05% Tween, PBS-T), blocked with blocking buffer for 1 hr (PBS-T with 5% BSA) and incubated with the pSVV antibody for 1 hr at room temperature. Next, wells were incubated with anti-rabbit antibody conjugated to Biotin and further incubated with Streptavidin-HRP for detection. For signal detection, TMB (1-Step Ultra TMB-ELISA, Thermo Scientific) was added, incubated for color development and stopped by addition of STOP solution (Thermo Scientific), which was read at 450 nm using an xMark Spectrophotometer plate reader (BioRad). Each sample was analyzed in triplicate.

For autophosphorylation and subsequent analysis by Western blot, 75 ng of purified CK1δ1-FL protein (from Signalchem) was incubated in kinase buffer (25 mM Tris pH 7.5, 5 mM beta glycerol phosphate, 2 mM DTT and 0.1 mM sodium orthovanadate) without or with 10 mM magnesium chloride and 200 μM ATP (for autophosphorylation) for 30 min at 25°C. Then, samples were added with SDS Buffer and processed according to the SDS-PAGE and Western blot protocol. Either purified protein samples or whole cell extracts of transfected HEK293 cells lysed with cell lysis buffer (50 mM Tris-HCl pH 8.0, 150 mM NaCl, 1% Nonidet P-40, and 0.5% deoxycholic acid containing Complete protease inhibitors (Roche) and PhosStop phosphatase inhibitors (Roche)) were analyzed by denaturing SDS-PAGE gel, which was transferred on PVDF membrane (Immobilon, Millipore). Blot was further probed using indicated primary antibodies and appropriate secondary antibodies conjugated with HRP. Signal was detected with ECL reagents from Thermo Fisher Scientific. Densitometric analysis of WB bands were performed using the ImageJ software (National Institutes of Health). The ratio of pS659 to myc-PER2 was calculated and values are expressed in Fig. 5G.

To detect autophosphorylation of CK1δ1 tail region, HEK293T cells were transfected with indicated expression plasmids and 48 hr after transfection cells were treated with either DMSO or 40 nM of Calyculin A (Cell Signaling Technology) for 1 hr and lysed in cell lysis buffer. For CK1 inhibition studies, before Calyculin A treatment cells were pretreated with either DMSO or 1 µM PF670 (PF670462, Tocris Bioscience) for the indicated duration. These samples were subsequently analyzed by Western blot and probed with pSVV and other indicated antibodies.

### Generation of stable PER2::LUC CK1δ lines

CK1δ1 WT or CK1δ1 S3A were synthesized and cloned into the mammalian lentiviral expression backbone (Addgene #73320) with a modification to include a C-terminal FLAG tag and a stop codon in-frame with the EGFP to prevent expression of the fusion protein (Twist Biosciences). Recombinant lentiviral particles were produced in HEK293T cells (ATCC) using Pax2 and pMD2.5 packaging plasmids. The resulting supernatant was used to transduce U2OS PER2::LUC fibroblasts. For selection, 1 μg/ml puromycin was applied for one week with medium changes every 48 h. Successfully transduced cells were grown to confluence in 6-well dishes in high-glucose (27.8 mM), glutamax-containing DMEM (GIBCO) supplemented with 10% serum (HyClone FetalClone III, Thermo Fisher Scientific) and penicillin–streptomycin. Reconstituted lines also had 0.5 μg/ml puromycin to maintain selection. Confluent cultures were kept for up to 4 weeks with the medium refreshed every 7–10 days. Before the start of recording, cells were synchronized by the addition of 100 nM dexamethasome for 1 h and then changed to MOPS-buffered ‘air medium’ (bicarbonate-free DMEM, 5 mg/ml glucose, 0.35 mg/ml sodium bicarbonate, 0.02 M MOPS, 100 μg/ml penicillin–streptomycin, 1% Glutamax, 1 mM luciferin, pH 7.4, 325 mOsm (64). Cells were then transferred to an Alligator system (Cairn Research), in which bioluminescent activity was recorded at 15-min intervals using an electron multiplying charge-coupled device (EM-CCD) at constant 37 °C. Bioluminescent traces of cells were fitted with damped cosine waves using the following equation:

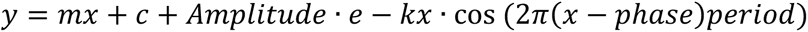

where *y* is the signal, *m* is the gradient of the detrending line, *c* is the *y* intercept of this detrending line, *x* is the corresponding time, amplitude is the height of the peak of the waveform above the trend line, *k* is the decay constant (such that 1/*k* is the half-life), phase is the shift relative to a cos wave and the period is the time taken for a complete cycle to occur.

### NMR spectroscopy

NMR spectra were collected on a Varian INOVA 600 MHz, or a Bruker 800 MHz spectrometer equipped with a ^1^H, ^13^C, ^15^N triple resonance z-axis pulsed-field-gradient cryoprobe. Spectra were processed using NMRPipe (65) and analyzed using CCPNmr Analysis (66). Spectra were collected in NMR buffer (25 mM MOPS pH 7.0, 1 mM EDTA, 100 mM NaCl, 5 mM BME, 11 mM MgCl_2_, and 0.05% Tween) with 10% D_2_O at 25°C. The backbone assignment of the δ1 and δ2 tails was accomplished using standard NH-edited triple-resonance experiments [HN(CA)CO, HNCO, HNCACB, and CBCA(CO)NH] and four-dimensional carbon direct-detection methods such as (HACA)N(CA)CON and (HACA)N(CA)NCO. Phosphopeak assignments of the CK1δ1 short tail were accomplished using standard NH-edited triple-resonance experiments [HN(CA)CO, HNCO, and HNCACB].

Chemical shift perturbation (Δδ) or the change in ^13^C-^15^N CON peak position was calculated using the following equation:

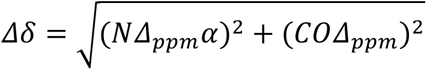

where Δ_ppm_ is the change in chemical shift and a value of α = 0.3 was used to normalize between ^15^N and ^13^C chemical shift ranges. NMR kinase reactions were performed at 25°C with 100 µM ^13^C,^15^N CK1δ1 tail or CK1δ2 tail, 2.5 mM ATP and 100 µM CK1δ ΔC. Samples were incubated for 2 hours and quenched with 20 mM EDTA, then ^1^H-^15^N HSQC spectra (total data acquisition = 20 min) and ^13^C-^15^N CON spectra (total acquisition time= 9 hr 32 min) were collected. The incubation time to reach saturation of the kinase reaction with CK1δ1 short tail was determined by collecting a series of ^1^H-^15^N HSQC spectra (53 20-min spectra for a total acquisition time = 1,060 min) of 100 µM ^13^C,^15^N CK1δ1 short tail, 2.5 mM ATP and 100 µM CK1δ ΔC at 25°C. For phosphopeak assignments, the CK1δ1 short tail kinase reaction was then performed with 100 µM ^13^C,^15^N CK1δ1 short tail, 2.5 mM ATP and 100 µM CK1δ ΔC incubated overnight at 25°C and quenched with 20 mM EDTA, then ^1^H-^15^N HSQC spectra (total data acquisition = 20 min), ^13^C-^15^N CON spectra (total acquisition time= 9 hr 32 min), and triple-resonance experiments were collected.

Segmentally labeled NMR kinase reactions were performed at 25°C with 34 μM CK1δ ΔC*-^15^N CK1δ1 and 2.5 mM ATP. Samples were prepared as described above, and then ^1^H-^15^N HSQC spectra (total data acquisition= 2 hours) were collected. For the phosphorylation kinetic assays, reactions were performed at 25°C with 34 µM CK1δ ΔC*-^15^N CK1δ1 tail, or 34 µM CK1δ ΔC* and 34 µM ^15^N CK1δ1 tail in *trans*, and then ^15^N-edited 1D ^1^H spectra were collected (224 data points at 6 min per spectrum for a total data acquisition= 1,344 min) immediately after the addition of 2.5 mM ATP. Spectra were processed using MestreNova (MestreLab), and data analysis was performed using Excel (Microsoft) and Prism (GraphPad).

### Hydrogen-deuterium exchange mass spectrometry

Sample preparation

To determine the effects of the nucleotide binding and monitor catalytic activity, hydrogen-deuterium exchange mass spectrometry (HDX-MS) was performed for different CK1δ constructs in the presence and absence of ATP or AMPPNP. Different CK1δ constructs were incubated with either ATP (∼800 μM) or AMPPNP (∼1.2 mM), or neither (apo), and 2 mM MgCl_2_ for 15 min at 25 °C before each hydrogen-deuterium exchange reaction. For HDX of CK1δ ΔC bound to the PER2 4p-FASP peptide, an excess of the peptide substrate (2 mM final concentration) was incubated with CK1δ ΔC in the presence of ADP and MgCl_2_. Hydrogen-deuterium labeling was initiated by diluting 60-65 pmol of CK1δ saturated with ATP or AMPPNP as indicated above in a reaction buffer prepared in deuterium oxide (D_2_O, final concentration of 90%). Pulse-labeling reactions were carried out for 1, 5, and 10-minute timepoints, after which the exchange was minimized by lowering the pH to 2.5 and temperature to 0°C by adding pre-chilled quench solution (0.5 M Guanidinium hydrochloride, 0.1 M TCEP). Non-deuterated controls were carried out by diluting nucleotide-free CK1δ proteins in an aqueous reaction buffer instead of a deuterated buffer.

Mass spectrometry data acquisition

For nanoLC-MS/MS analyses, each quenched sample was subjected to in-line proteolysis using the Enzymate^TM^ immobilized pepsin cartridge (Waters, USA). The digestion was left to proceed for 3 minutes at 12°C, and the resulting peptide mixtures were analyzed with nanoACQUITY M-class UPLC (Waters, UK) equipped with a trapping column (VanGuard C18, 5 μm particle size) and an analytical column (BEH C18, 75 μm × 100 mm, 1.7 μm particle size) maintained at 3°C to minimize back-exchange (67). The aqueous solvent was 0.1% formic acid in LC-MS grade water, and the organic solvent was 0.1% formic acid in HPLC-grade acetonitrile. A 10-minute gradient was used to resolve and elute the peptides by reverse-phase chromatography. The peptides were then identified by Synapt G2-Si high-resolution mass spectrometer (Waters, UK). Briefly, the mass spectrometer was operated in positive polarity, data-independent (DIA) fragmentation and ion-mobility (HDMS^E^) mode separation. Mass spectrometer parameters were as described previously (68). Glu-fibrinopeptide reference (m/z = 785.8426) was continuously sprayed every 30 seconds during ESI-MS experiments using the lockspray device to ensure mass accuracy.

HDX data analysis

Peak lists for identifying and annotating the peptides from non-deuterated controls were generated by Protein Lynx Global Server v3.0 (Waters, USA), using human CK1δ (Uniprot: P48730) construct-specific protein database and searched against the parameters described previously (68). Peptide mass measurements for individual peptides were assigned using DynamX v3.0 software (Waters, USA). Peptides with a minimum intensity of 2000, maximum length of 25 amino acids, and mass error within ± 10 ppm were considered for analysis. Deuterium exchange measurements were determined for all labeling timepoints across different protein states and were manually verified, but the data were not corrected for back-exchange. Only peptides with high signal-to-noise ratio and non-overlapping spectra were considered for final analysis. All measurements were done in biological duplicates and technical triplicates, with statistical analysis done using Deuteros 2.0 (69) and tabulated as Supplementary Tables 1-3.

HDX data representation

Relative fractional deuterium uptake (RFU) is the ratio of the number of deuterons exchanged to the maximum number of available amides per peptide. Differences in deuterium exchange (ΔDU) were calculated as differences in centroid values of a deuterated peptide in protein condition with its corresponding deuterated peptide in a different protein condition. The pepsin digestion maps showing the sequence coverage of various CK1δ constructs and their corresponding deuterium uptake values are provided in Supplementary Tables 1 and 2. HDX profiles between different nucleotide bound CK1δ constructs indicated in Supplementary Figure S4 were compared by subtracting the RFU values for common peptides, as per the following scheme:

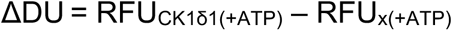

where x = CK1δ2, CK1δ1 S3A, CK1δ Δ400, or CK1δ ΔC

### QUANTIFICATION AND STATISTICAL ANALYSIS

All statistical analyses were done using Prism (GraphPad). P-values were calculated using Students t-test, one-way ANOVA with Tukey’s multiple comparisons test, or extra sum of squares F-test as indicated in different figures. In all figures, * indicates p<0.05, **p<0.01, ***p<0.001, ****p<0.0001, ns, not significant.

## Acknowledgements

Funding for this work was provided by the US National Institutes of Health grants R01 GM107069 and R35 GM141849 (to C.L.P.), and Singapore Ministry of Health grant MOH-000600 to D.M.V. We thank Andrew D. Beale and John S. O’Neill (MRC LMB, UK) for their generous gift of the U2OS PER2::LUC cell line, Jae Kyoung Kim (KAIST, Korea) for consultations on the mathematical model.

## Author contributions

Conceptualization: R.H., N.K.T., R.N., C.L.P., and D.M.V.; Investigation: R.H., N.K.T., R.N., N.Y., M.A.H., H.-W.L., P.C., and S.T.; Formal analysis: R.H., N.K.T., R.N., N.Y., H.-W.L., P.C., and S.T.; Writing – Original draft: R.H., N.K.T., R.N., and C.L.P.; Writing – Review & Editing: R.H., N.K.T., R.N., C.L.P., and D.M.V.; Supervision: C.L.P. and D.M.V.; Funding Acquisition: C.L.P. and D.M.V.

## Supplementary Material

**Supplementary Table 1**

Deuterium exchange values for various pepsin-digested peptides of CK1δΔC in the presence of ATP, AMPPNP, and 4p-FASP at 1, 5, and 10-minute labeling times are tabulated.

**Supplementary Table 2**

Deuterium exchange values for various pepsin-digested peptides of CK1δ1, CK1δ2, CK1δ1 S3A, and CK1δ1ΔC in the presence of ATP, AMPPNP, and 4p-FASP at 1, 5, and 10-minute labeling times are tabulated.

**Supplementary Table 3**

HDX metadata table depicting the experimental conditions for different CK1δ1 isoforms, and the corresponding analysis is represented.

**Supplementary Table 4**

X-ray crystallography data collection and refinement statistics.

