## Supplemental figures for "Isoform-specific C-terminal phosphorylation drives autoinhibition of Casein Kinase 1"

Sup Fig 1

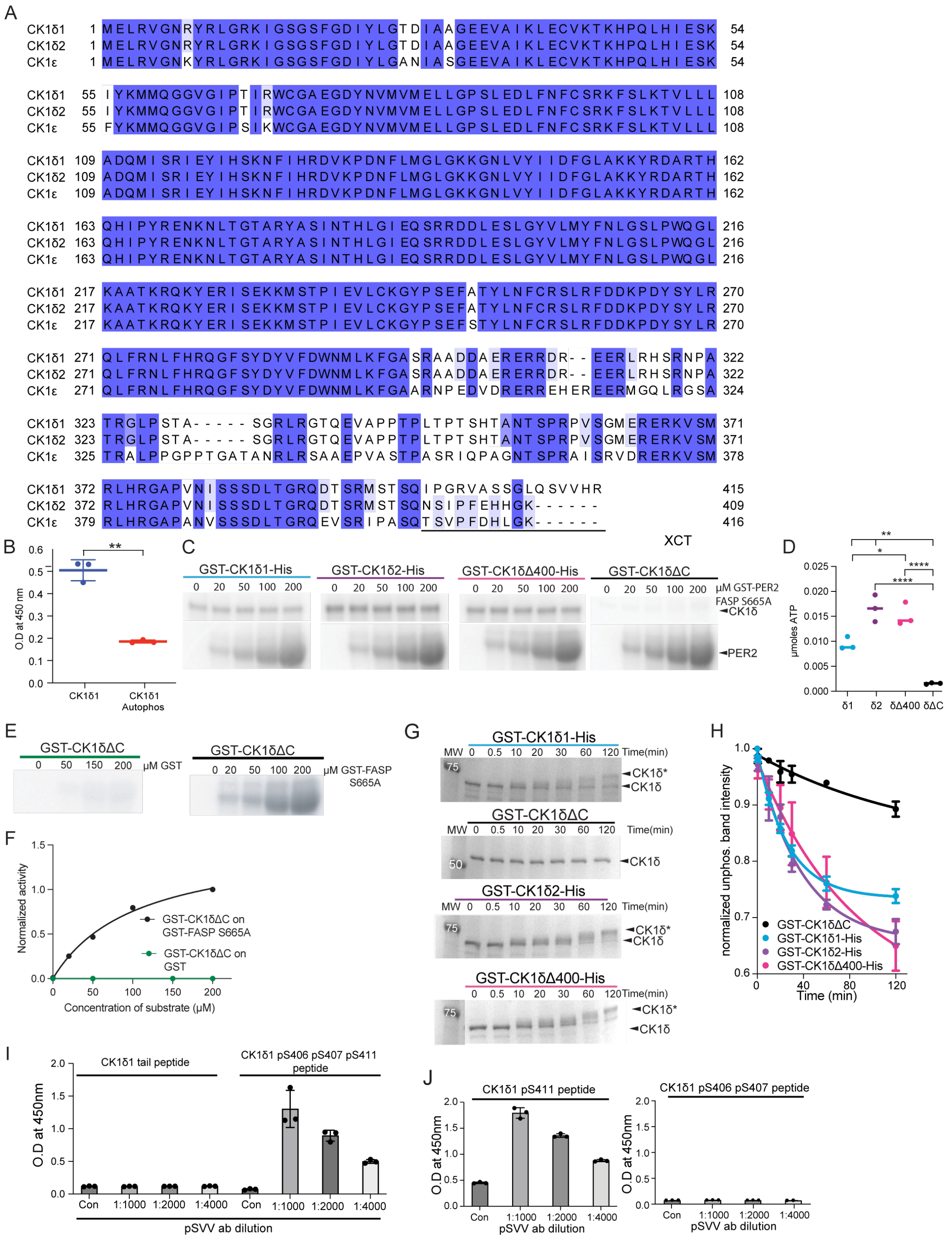


### Supplemental Figure 1

**A**, Sequence alignment of human CK1δ isoforms δ1 and δ2 with CK1ε. Kinase domain, residues 1-294. **B**, ELISA-based peptide kinase assay showing phosphorylation of the mouse PER2 FASP priming serine (pS659) with 50 ng CK1δ1 after pretreatment without (blue) or with (red) ATP to allow for kinase autophosphorylation. (n=3 mean ± s.d.). **C,** Autoradiograph of $^{32}P$ kinase assay; top, autophosphorylation of kinase; bottom, priming phosphorylation of the PER2 FASP substrate. Representative autoradiograph from n=4. **D**, Quantification of autophosphorylation in panel C. Ordinary one-way ANOVA, * p>0.05; **, p<0.01; ****, p<0.0001 (n=4, mean ± s.d.). **E**, Representative autoradiograph of $^{32}P$ kinase assay of CK1δΔC activity on GST-human PER2 FASP S665A or GST alone (n=1). **F**, Quantification of phosphorylation in panel E. **G,** Representative Coomassie SDS-PAGE gel of autophosphorylation time course with CK1δ1 (blue), CK1δΔC (black), CK1δ2 (purple), or CK1δΔ400 (pink) n=3. Hyperphosphorylated kinase indicated by CK1δ*. **F,** Quantification of panel G for the unphosphorylated band intensity over time for CK1δ1 (blue), CK1δΔC (black), CK1δ2 (purple), or CK1δΔ400 (pink) normalized to time 0 band intensity (n=3, mean +/- SEM). **H,** ELISA of unphosphorylated CK1δ1 tail peptide or a triply phosphorylated CK1δ1 tail peptide with the pSVV ab (n=3 mean ± s.d.). **I,** ELISA of CK1δ1 pS411 tail peptide or CK1δ1 pS406 pS407 tail peptide with the pSVV ab (n=3 mean ± s.d.).

Sup Fig 2

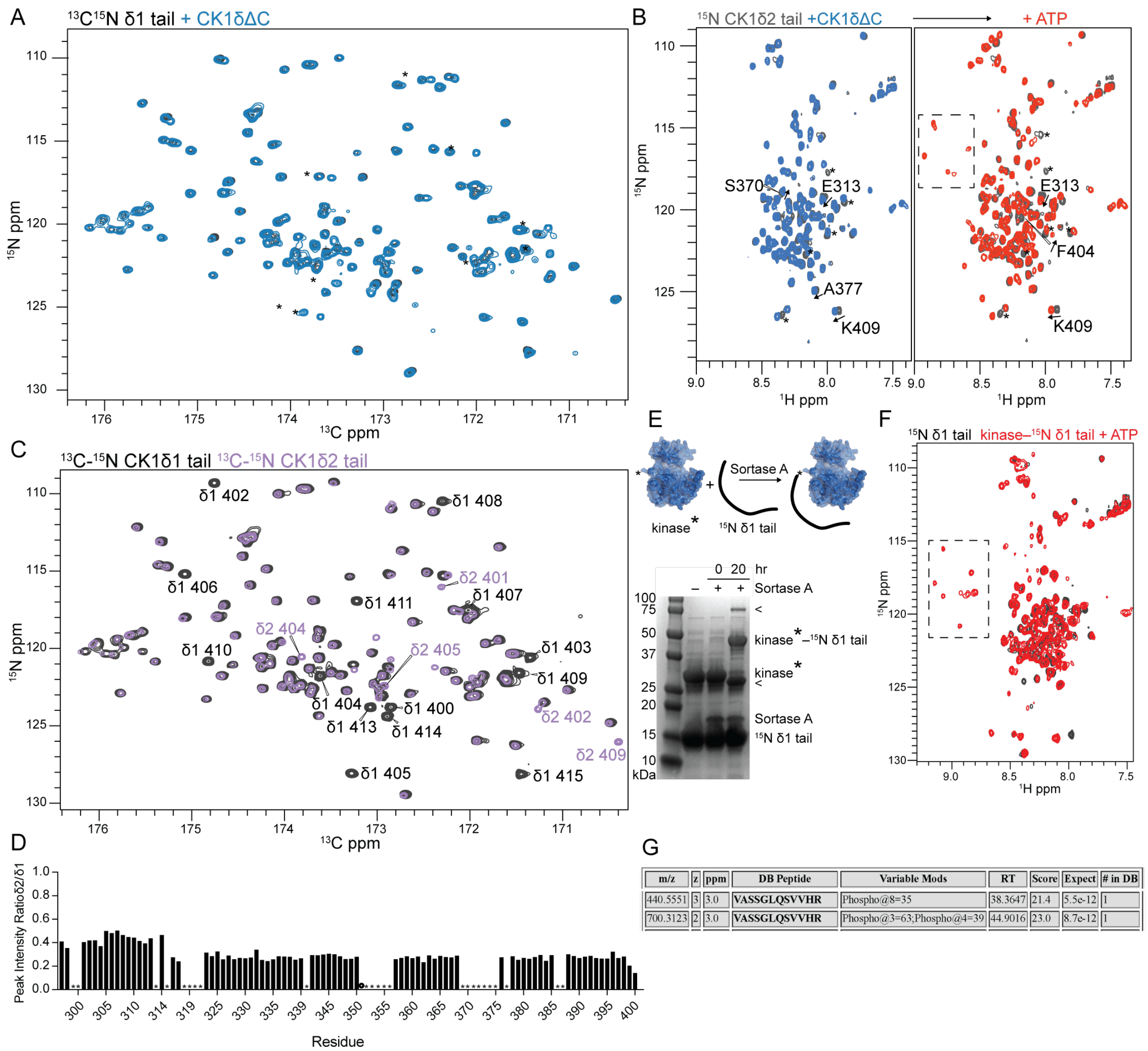


| m/z | z | ppm | DB Peptide | Variable Mods | RT | Score | Expect | # in DB |
|---|---|---|---|---|---|---|---|---|
| 440.5551 | 3 | 3.0 | **VASSGLQSVVHR** | Phospho@8=35 | 38.3647 | 21.4 | 5.5e-12 | 1 |
| 700.3123 | 2 | 3.0 | **VASSGLQSVVHR** | Phospho@3=63;Phospho@4=39 | 44.9016 | 23.0 | 8.7e-12 | 1 |

**Supplemental Figure 2**

**A**, $^{15}$N-$^{13}$C CON spectra of 100 μM $^{13}$C,$^{15}$N δ1 tail alone (black) or in the presence of 100 μM CK1δΔC without ATP (blue). **B**, $^{15}$N-$^{1}$H HSQC spectra of 100 μM $^{15}$N δ2 tail alone (black) or in the presence of 100 μM CK1δΔC (blue) and 2.5 mM ATP (red) for 2 hours at 25 ºC; dashed box, peaks corresponding to phosphorylated residues. Arrows, chemical shift perturbation with kinase alone or with phosphorylation. **C**, $^{15}$N-$^{13}$C CON spectra of 400 μM $^{13}$C,$^{15}$N δ1 tail alone (black) compared to 400 μM $^{13}$C,$^{15}$N δ2 tail alone (purple) with isoform-specific assignments. **D**, Peak intensity of $^{15}$N-$^{13}$C CON δ2 compared to δ1; *, unassigned peaks. **E**, Synthetic scheme for segmental isotopic labeling of the δ1 tail in a full-length kinase. **F**, $^{15}$N-$^{1}$H HSQC spectra of 34 μM $^{15}$N δ1 tail alone (black) or the segmentally labeled kinase with 2.5 mM ATP (red) for 2 hours at 25ºC. **G,** Identification of δ1-specific phosphorylation sites from full-length autophosphorylated CK1δ1 after trypsin digest and LC/MS-MS.

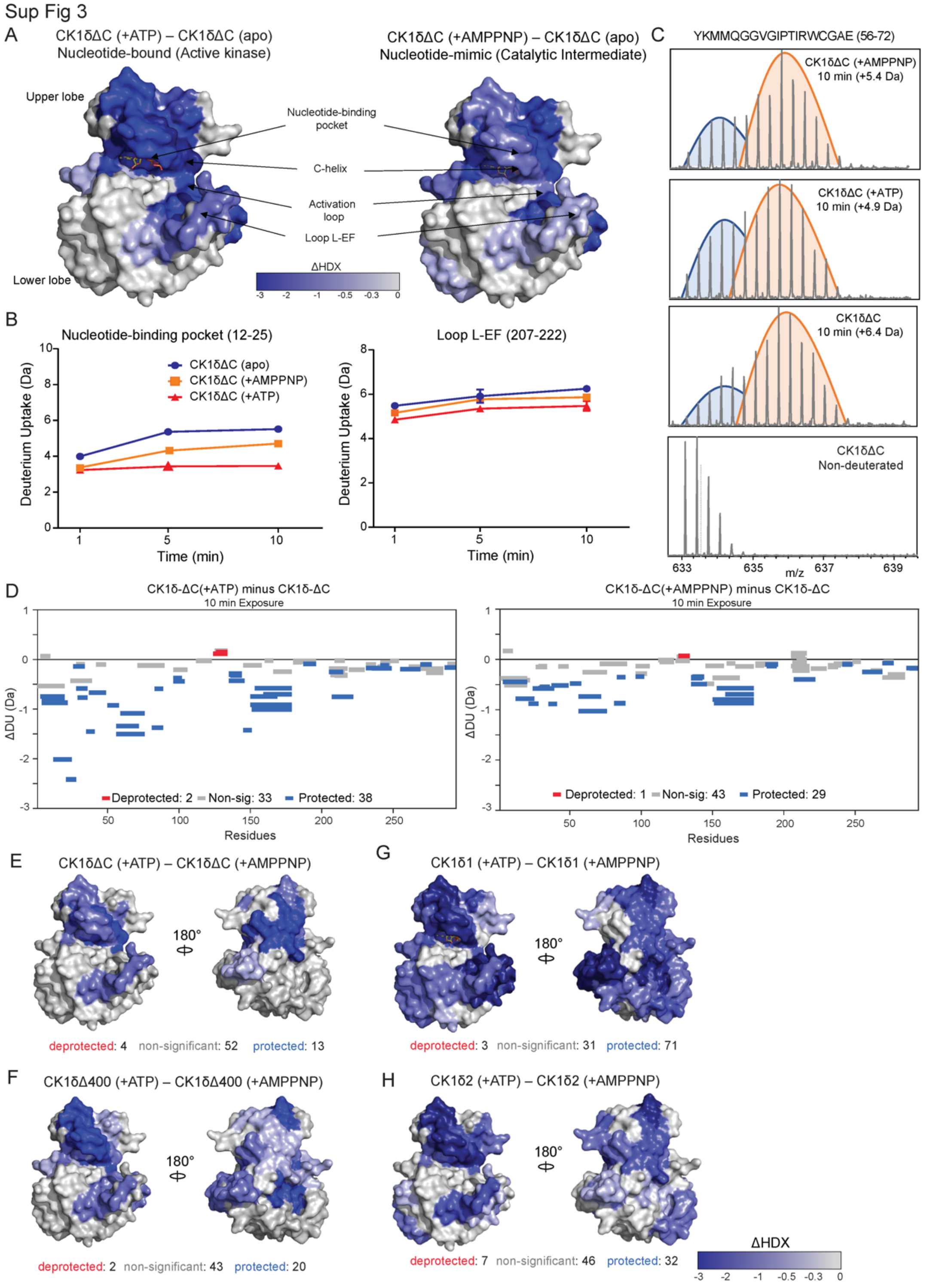
Sup Fig 3
A
CK1δΔC (+ATP) – CK1δΔC (apo)
Nucleotide-bound (Active kinase)
CK1δΔC (+AMPPNP) – CK1δΔC (apo)
Nucleotide-mimic (Catalytic Intermediate)
Upper lobe
Lower lobe
Nucleotide-binding pocket
C-helix
Activation loop
Loop L-EF
ΔHDX
-3 -2 -1 -0.5 -0.3 0
B
Nucleotide-binding pocket (12-25)
Loop L-EF (207-222)
CK1δΔC (apo)
CK1δΔC (+AMPPNP)
CK1δΔC (+ATP)
Deuterium Uptake (Da)
Time (min)
C
YKMMQGGVGIPTIRWCGAE (56-72)
CK1δΔC (+AMPPNP) 10 min (+5.4 Da)
CK1δΔC (+ATP) 10 min (+4.9 Da)
CK1δΔC 10 min (+6.4 Da)
CK1δΔC Non-deuterated
m/z
D
CK1δ-ΔC(+ATP) minus CK1δ-ΔC
10 min Exposure
Deprotected: 2 Non-sig: 33 Protected: 38
CK1δ-ΔC(+AMPPNP) minus CK1δ-ΔC
10 min Exposure
Deprotected: 1 Non-sig: 43 Protected: 29
ΔDU (Da)
Residues
E
CK1δΔC (+ATP) – CK1δΔC (+AMPPNP)
180°
deprotected: 4 non-significant: 52 protected: 13
F
CK1δΔ400 (+ATP) – CK1δΔ400 (+AMPPNP)
deprotected: 2 non-significant: 43 protected: 20
G
CK1δ1 (+ATP) – CK1δ1 (+AMPPNP)
deprotected: 3 non-significant: 31 protected: 71
H
CK1δ2 (+ATP) – CK1δ2 (+AMPPNP)
deprotected: 7 non-significant: 46 protected: 32
ΔHDX
-3 -2 -1 -0.5 -0.3 0

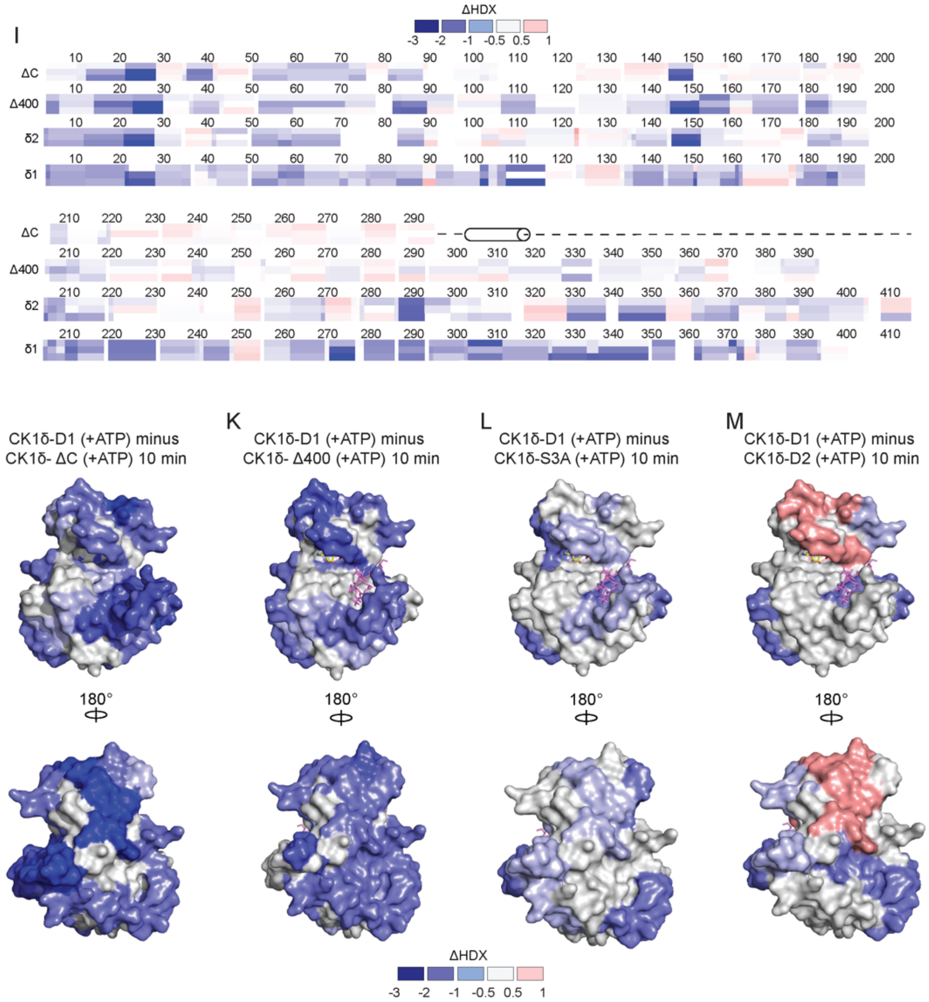


**Supplemental Figure 3 Conformational changes mediated by nucleotide binding to CK1δ**

**A**, Structural representations of CK1δ kinase domain (PDB: 6PXO, with ATP (yellow) from PDB: 6RU6) indicating the regions with reduced deuterium exchange when saturated with ATP (left) or the ATP-analogue AMPPNP (right). Differences in deuterium exchange at 10 min labeling time are mapped in shades of blue as indicated. **B,** Kinetic plot of relative deuterium uptake for peptides spanning the nucleotide-binding pocket (left) and the catalytic loop (right) for CK1δΔC apo (blue), with ATP (red), or AMPPNP (orange). Average values (n = 6) from technical and biological replicates were used to generate the plot. **C,** Stacked mass spectral plots for peptide containing residues 56-72 with bimodal isotopic distribution at 10 min labeling time with lower (blue) and higher (orange) exchanging populations of CK1δΔC apo, or with ATP or AMPPNP, and a non-deuterated control. **D,** Woods differential plots showing the effect of ATP (top) or AMPPNP (bottom) relative to apo CK1δΔC. Plots generated using Deuteros 2.0. The 99.0% confidence interval was used to highlight peptides with significant deprotection (red), or protection (blue) compared to no change (gray) in deuterium exchange. **E-H,** structural representation of deuterium exchange differences at 10 min corresponding to Fig 3 E-F with number of peptides deprotected (red), non-significant (gray), and protected (blue) below each structure. **I,** Heat map of changes by residue in relative fractional uptake at 1 (top), 5 (middle), and 10 (bottom)-min labeling times for indicated constructs between ATP-bound and AMPPNP-bound states. **J-M,** Structural representations of the differences in relative deuterium uptake values between CK1δ1 with different isoforms **J**, CK1δΔC, **K**, CK1δΔ400, **L**, CK1δ1 S3A, and **M**, CK1δ2 in ATP-bound states. The differences in RFU are mapped onto the kinase domain (PDB: 6PXO) in front (top) and rear (bottom) views and are calculated as described in methods.

Sup Fig 4

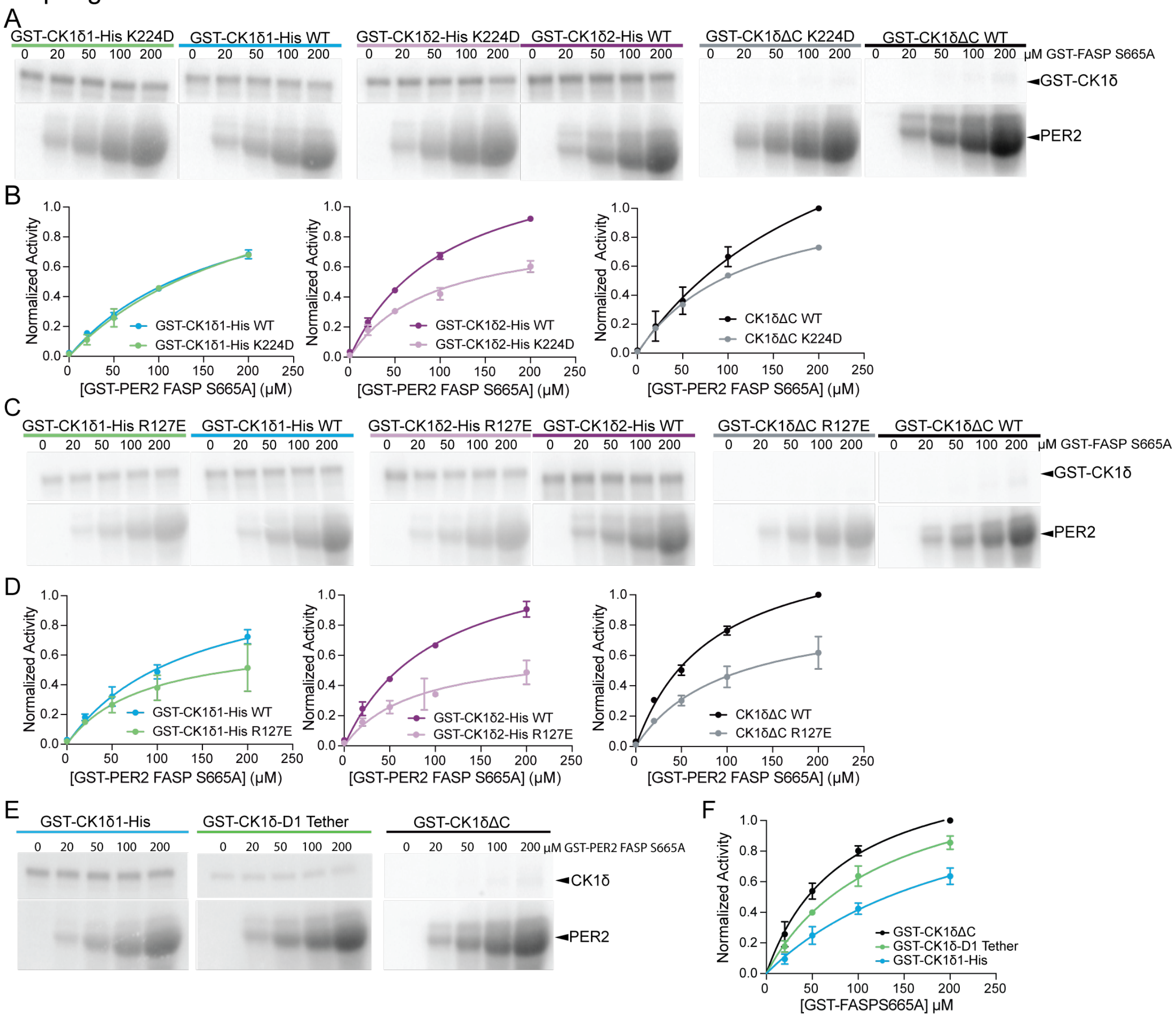


#### Supplemental Figure 4

**A**, Representative autoradiographs from $^{32}$P kinase assays of Site 1 mutant K224D or WT kinase construct as indicated on GST-PER2 FASP S665A (n=3). **B**, $^{32}$P kinase assays of anion binding site 1 mutant, K224D, or WT in CK1δΔC (gray/black), CK1δ1 (green/blue), and CK1δ2 (lavender/purple) on PER2 GST-FASP-S665A (n=3, mean ± s.d.). **C**, Representative autoradiographs from $^{32}P$ kinase assay of Site 2 mutant R127E or WT kinase construct as indicated on GST-PER2 FASP S665A (n=3). **D**, $^{32}P$ kinase assays comparing activity of anion binding site 2 mutant, R127E, or WT in CK1δ-ΔC (gray/black), CK1δ1 (green/blue), and CK1δ2 (lavender/purple) on PER2 GST-FASP-S665A (n=3, mean ± s.d.). **E,** Representative autoradiographs from $^{32}P$ kinase assays of CK1δ1 (blue), CK1δ-D1 tether (green), and CK1δΔC (black) on GST-PER2 FASP S665A (n=3). **F,** Quantification of $^{32}P$ kinase assays of CK1δ1 (blue), CK1δ-D1 tether (green), and CK1δΔC (black) on GST-PER2 FASP S665A (n=3, mean +/- s.d.).

Sup Fig 5

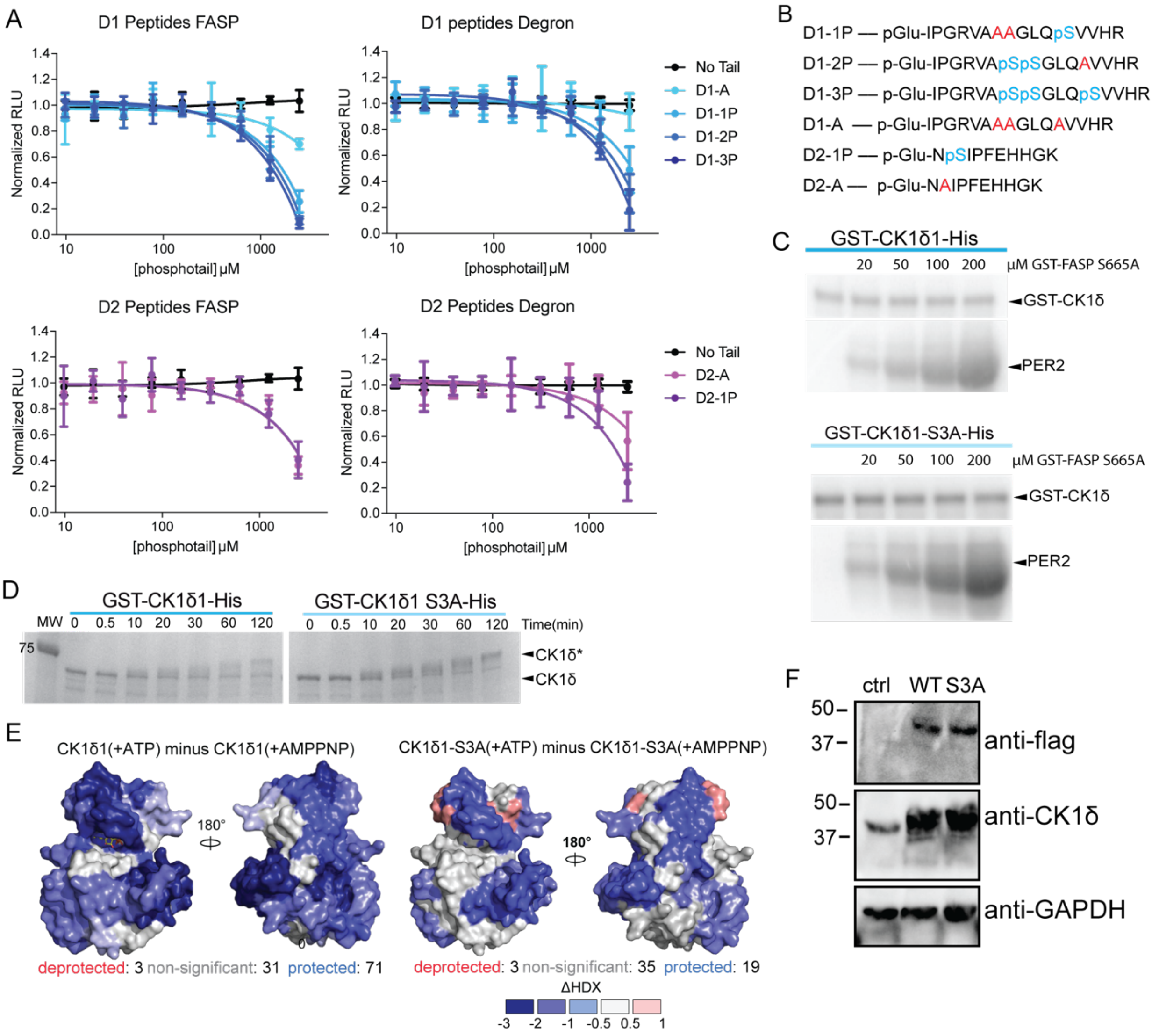


**Supplemental Figure 5**

**A**, ADP-Glo kinase assay with phosphorylated CK1δ C-terminal peptides, D1 peptides (blue) and D2 peptides (purple), on PER2 FASP (left) or Degron (right). (n=2) **B,**

Phosphorylated CK1δ C-terminal peptides; blue, phosphorylated serines (pS); red, alanine mutations to block phosphorylation. **C,** Autoradiograph of $^{32}P$ kinase assay measuring autophosphorylation of kinase (top) or priming phosphorylation of the PER2 FASP substrate (bottom) for CK1δ1 (blue) and CK1δ1 S3A (light blue). Representative autoradiograph from n = 3. **D,** Representative coomassie SDS-PAGE gel of autophosphorylation time course with CK1δ1 (blue) and CK1δ1 S3A (light blue) n=3. Hyperphosphorylated kinase indicated by CK1δ*. **E,** Structural representation of the CK1δ kinase domain (PDB: 6PXO, with ATP (yellow) from PDB: 6RU6) indicating the regions with differing deuterium exchange in CK1δ1 (top) and CK1δ1 S3A (bottom) comparing ATP to AMPPNP-bound states (right). Differences in deuterium exchange at 10 min labeling time are mapped in shades of blue as indicated. Number of peptides deprotected (red), non-significant (gray), and protected (blue) indicated below each structure. **F,** Western of stably expressed FLAG-tagged CK1δ1 or CK1δ1 S3A in PER2::LUC U2OS cells compared to endogenous expression of CK1δ in untreated control PER2:LUC U2OS cells.
